# Astrocytic glycolysis attenuates mitochondrial efficiency to preserve cognition

**DOI:** 10.64898/2025.12.24.696355

**Authors:** Paula Alonso-Batan, Daniel Jimenez-Blasco, Jesus Agulla, Rebeca Lapresa, Marta Antequera-Düwel, Sara Yunta-Sanchez, Brenda Morant-Ferrando, Dario Garcia-Rodriguez, Rebeca Acin-Perez, Veronica Bobo-Jimenez, Leticia Sancha-Ortega, Regina Mengual, Silvia Gomila, Sonia Gonzalez-Guerrero, Emilio Fernandez, Gilles Bonvento, Jose A. Enriquez, Peter Carmeliet, Angeles Almeida, Juan P. Bolaños

## Abstract

Astrocytic glycolysis is tightly coupled to neurotransmission and thought to be essential for neurological health. However, the metabolic adaptations that enable astrocytes to maintain a durable glycolytic profile without compromising viability are elusive. Here, using *in vivo* approaches including cell-specific gene expression disruption, metabolic flux analyses and behavioral tests in mice, we addressed this issue. We found that Pfkfb3 (6-phosphofructo-2-kinase/fructose-2,6-bisphosphatase-3) is instrumental in maintaining the astrocytic glycolytic phenotype. Importantly, astrocytic glycolysis sustained by Pfkfb3 is required for normal cognitive performance. Mechanistically, ATP generated through glycolysis is consumed by mitochondria, *via* the reverse mode of ATP synthase, to conserve the proton gradient across the inner mitochondrial membrane. This enables mitochondria to attenuate pyruvate decarboxylation, tricarboxylic acid cycle and electron transport chain activity, thereby preserving pyruvate for conversion into lactate and delivery to neurons. These findings reveal that astrocytes sacrifice mitochondrial bioenergetic efficiency as a previously underappreciated strategy to support cognition.

## Main text

In contrast to neurons^1–3^, neighbors’ astrocytes robustly consume glucose through glycolysis^1,4–7^. It is thought that astrocytic glycolysis is key to drive neurotransmission by means of delivering metabolites, such as lactate and serine, which serve as neuronal fuels and signals^8–13^ to preserve neurological wellbeing^14,-16^. However, despite its critical importance, the molecular mechanisms that enable astrocytes to constantly dispense with oxidative fuel for neuronal support, while preserving viability remains unsolved. According to *in vitro* studies, Pfkfb3 -an enzyme that synthesizes the potent glycolytic activator fructose-2,6-bisphosphate (F26BP)^17^, evades proteasomal degradation in astrocytes thereby accounting for a high stability and enzyme activity in these cells^1^. This suggests that Pfkfb3 may be an excellent regulatory candidate responsible for the *in vivo* maintenance of a durable glycolytic flux in astrocytes to sustain brain function. We therefore aimed to address the impact of astrocytic glycolysis on organismal behavior by genetically engineering adult mice lacking *Pfkfb3* in astrocytes. This was achieved by promoting Cre recombinase-mediated deletion of the *Pfkfb3* gene fragment that flanked exons 3 to 6, selectively in astrocytes, using previously generated *Pfkfb3^lox/lox^*mice^18^. To induce *Pfkfb3* gene deletion *in vivo*, we used a previously validated strategy^19^ in which 3 months-old *Pfkfb3^lox/lox^* male mice were intravenously injected, *via* the retroorbital sinus^20^, with PHP.eB serotype adeno-associated virus particles expressing Cre recombinase governed by the astrocytic-specific glial-fibrillary acidic protein (GFAP) short-promoter (PHP.eB-AAV-gfaABC_1_D-Cre-GFP) (**Fig. 1a; Extended Data Fig. 1a**). Controls (wild type, WT) were *Pfkfb3^lox/lox^* mice that received equivalent doses of the same virus particles, except that they lacked Cre recombinase. After 2 months, Pfkfb3 protein levels were substantially decreased in brain hippocampal extracts (**Extended Data Fig. 1b**) and in immunomagnetically isolated astrocytes (**Fig. 1b**) from the brain of the astrocyte-specific *Pfkfb3* knockout (KO) mice without altering other Pfkfb isoforms (**Extended Data Fig. 1c**). This was accompanied by a ∼61% decline in the flux of glycolysis, as assessed by [3-^3^H]glucose incorporation into ^3^H_2_O, a *bona fide* measure of glycolysis, in astrocytes isolated from *Pfkfb3* KO mice (**Fig. 1c**).

**Fig. 1.**
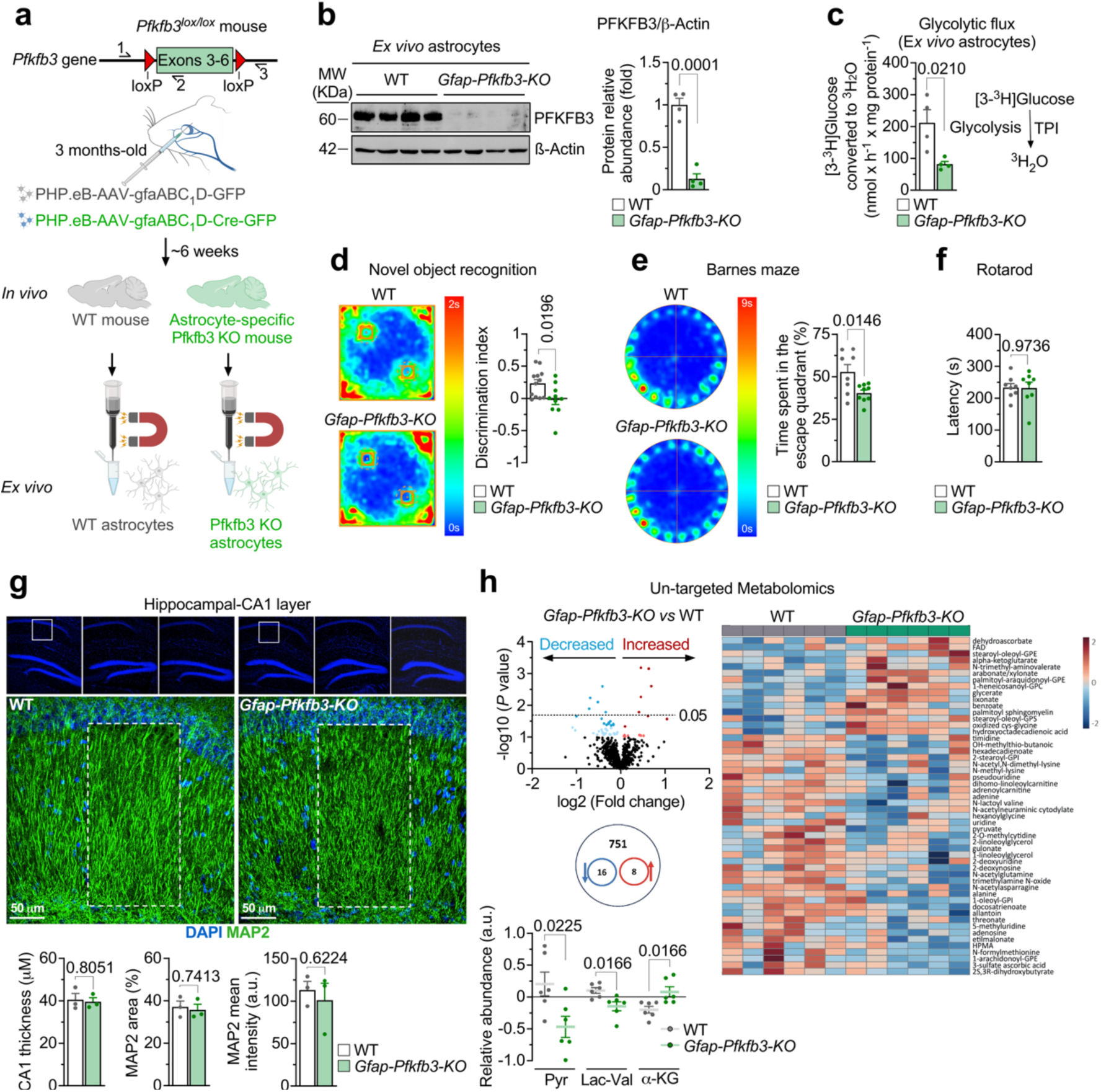
Astrocyte-specific repression of glycolysis by knocking out *Pfkfb3* impairs cognitive performance and alters brain metabolomic signature. **(a)** Strategy used to generate astrocyte-specific *Pfkfb3* knockout (*Gfap-Pfkfb3-KO*) mice and to immunomagnetically purify Pfkfb3-KO (and wild type, WT) astrocytes from adult brain. Created with BioRender.com. (See also Extended Data Fig. 1a). **(b)** Western blot against PFKFB3 protein in astrocytes immunomagnetically isolated from WT and *Gfap-Pfkfb3-KO* mouse brains; β-Actin was used as loading control (*left*). Quantification of PFKFB3 protein relative to β-Actin (*right*). Data are mean ± S.E.M. *P* value is indicated; n=4 mice per genotype; Unpaired Student’s t-test, two-tailed. (See also Extended Data Figs. 1b,c). **(c)** Glycolytic flux in astrocytes immunomagnetically isolated from WT and *Gfap-Pfkfb3-KO* mouse brains. Data are mean ± S.E.M. *P* value is indicated; n=4 mice per genotype; Unpaired Student’s t-test, two-tailed. (TPI, triosephosphate isomerase). **(d)** Novel object recognition test average occupancy heat maps in WT (*top*) and *Gfap-Pfkfb3-KO* (*bottom*) male mice (*left*). Discrimination index quantification (*right*). Data are mean ± S.E.M. *P* value is indicated; n=12 (WT) or 10 (*Gfap-Pfkfb3-KO*) mice; Unpaired Student’s t-test, two-tailed. (See also Extended Data Figs. 1d-j). **(e)** Barnes maze test average occupancy heat maps in WT (*top*) and *Gfap-Pfkfb3-KO* (*bottom*) mice (*left*). Quantification of the time spent in escape target quadrant (*right*). Data are mean ± S.E.M. *P* value is indicated; n=8 (WT) or 9 (*Gfap-Pfkfb3-KO*) mice; Unpaired Student’s t-test, two-tailed. **(f)** Rotarod test in WT and *Gfap-Pfkfb3-KO* mice. Data are mean ± S.E.M. *P* value is indicated; n=8 mice per genotype; Unpaired Student’s t-test, two-tailed. **(g)** Hippocampal CA1 structural analysis. *Upper panels*: DAPI- and MAP2-stained hippocampal sections from WT (*left*) and *Gfap-Pfkfb3-KO* (*right*) mice; white boxes highlight CA1 region shown in magnified panels below. Scale bar: 50 μm. *Lower panels*: quantification of CA1 thickness, MAP2-positive area, and mean MAP2 intensity of the dashed white boxes. Data are mean ± S.E.M. *P* values are indicated; n=3 mice per genotype; Unpaired Student’s t-test, two-tailed. (See also Extended Data Figs. 1k,l). |**(h)** Untargeted metabolomics of brain samples comparing *Gfap-Pfkfb3-KO versus* WT: *Left-top:* volcano plot showing metabolites significantly decreased (blue) or increased (red); horizontal dashed line indicates *P*=0.05 threshold. *Left-middle*: Venn diagram indicating the number of significantly altered metabolites (blue arrow decreases, red increases; total detected metabolites noted). *Right:* heat map of relative abundance of identified metabolites across WT and *Gfap-Pfkfb3-KO* samples, log₂ fold-change color scale. *Below*: bar graphs of relative abundances for key altered metabolites (pyruvate Pyr; lactate–valine, Lac–Val; and α-ketoglutarate, α-KG). Data are mean ± S.E.M. *P* values are indicated; n=3 mice per genotype; Welch’s t test. (See also Extended Data Fig. 1m).

Behavioral analysis in *Pfkfb3* KO mice revealed deficits in the short-term and spatial memories, as judged by the impaired performance in the novel object recognition test (**Fig. 1d**) and Barnes maze (**Fig. 1e**), respectively. However, locomotor activity was unaltered (**Fig. 1f**). According to the open field (**Extended Data Fig. 1d**) and the black and white box (**Extended Data Fig. 1e**) tests, astrocyte-specific *Pfkfb3* KO mice showed anxiety-like behavior, but the exploration capacity and the working memory were unaltered, as judged by the hole board (**Extended Data Fig. 1f**) and Y-maze (**Extended Data Fig. 1g**) tests, respectively. Female *Pfkfb3* KO mice showed unaltered behavioral and cognitive performances (**Extended Data Fig. 1h-j**). Immunocytochemical analysis showed unaltered neuronal morphology (**Fig. 1g**) and reduced astrocytic marker GFAP immunofluorescence intensity and roundness (**Extended Data Fig. 1k**), without changes in the proportion of Iba1^+^ (microglia) (**Extended Data Fig. 1l**) cells.

Next, we aimed to ascertain the metabolic alterations caused in the brain by the reduction of glycolysis in astrocytes. Brain (mostly cortex and hippocampus) untargeted metabolomics (**Fig. 1h**, *top panel***; Extended Data Fig. 1m**) revealed 16 metabolites significantly decreased and 8 metabolites significantly increased in the brain of the astrocyte-specific *Pfkfb3* KO mice (see **Source Data**). A closer analysis of the altered metabolites revealed decreased the glycolysis-end product pyruvate, and lactyl-valine - known to positively correlate with lactate abundance^21^ in the astrocytic-specific *Pfkfb3* knockout mice (**Fig. 1h**, *bottom panel*), in consonance with the observed reduction in the glycolytic flux (**Fig. 1c**). Interestingly, the mitochondrial tricarboxylic acid (TCA) cycle intermediate α-ketoglutarate increased (**Fig. 1h**, *bottom panel*). These data illustrate the efficacy of astrocyte-specific *Pfkfb3* KO strategy to downmodulating brain glycolysis *in vivo*, which may lead to enhanced mitochondrial TCA cycle activity.

To better characterize the metabolic alterations caused by the loss of *Pfkfb3*, astrocytes in primary culture obtained from *Pfkfb3^lox/lox^* mice were transduced with adenoviruses expressing Cre recombinase under the potent cytomegalovirus (CMV) promoter (AdV-CMV-Cre) (**Fig. 2a**). *Pfkfb3^lox/lox^*astrocytes transduced with the AdV lacking Cre recombinase (AdV-CMV-Ø) were used as controls (WT). As shown in **Fig. 2b**, Pfkfb3 protein was dose- and time-dependently decreased efficiently in AdV-CMV-Cre transduced astrocytes when compared with WT cells, without altering other Pfkfb isoforms (**Extended Data Fig. 2a**). Notably, the product of Pfkfb3 -and all Pfkfb isoforms- activity, F26BP, decreased by ∼50% (**Fig. 2c**), indicating that only Pfkfb3 contributes by at least half of the endogenous F26BP abundance in astrocytes. In good agreement with the *ex vivo* data (**Fig. 1c**), the glycolytic flux was reduced by ∼60% in Pfkfb3 KO primary astrocytes (**Fig. 2d**). To assess whether Pfkfb3 contributes to basal or stimulated glycolysis, astrocytes were challenged to a bioenergetic stress with the cytochrome *c* oxidase inhibitor, potassium cyanide (KCN) -known to promote 5’-AMP protein kinase (AMPK)-mediated Pfkfb2^22^ and Pfkfb3^6^ phosphorylation and activation. As shown in **Fig. 2d**, KCN enhanced by ∼3-fold the flux of glycolysis both in WT and *Pfkfb3* KO astrocytes, suggesting that, whilst Pfkfb3 accounts for basal glycolysis, other Pfkfb isoforms -likely, Pfkfb2- accounts for acute regulation. Notably, the changes observed in the glycolytic flux were replicated by analyzing the release of lactate (**Fig. 2e**). Given that astrocytic glycogen breakdown delivers lactate^23–25^, we assessed glycogen abundance. *Pfkfb3* loss resulted in glycogen accumulation and in strong impairment of glucose withdrawal-induced glycogen breakdown in astrocytes (**Fig. 2f**, *left panel*). However, glycogen was virtually exhausted by KCN treatment both in WT and *Pfkfb3* KO astrocytes (**Fig. 2f**, *right panel*). Of note, glucose metabolism through the pentose-phosphate pathway (PPP) was unaltered by *Pfkfb3* loss (**Fig. 2g; Extended Data Fig. 2b**), suggesting that unlike in neurons^1,2^, glycolysis and PPP are not reciprocally modulated in astrocytes. Altogether, these data strongly suggest that Pfkfb3-mediated glycolysis in astrocytes is necessary for sustained, rather than stimulated, glycogen mobilization and lactate release.

**Fig. 2.**
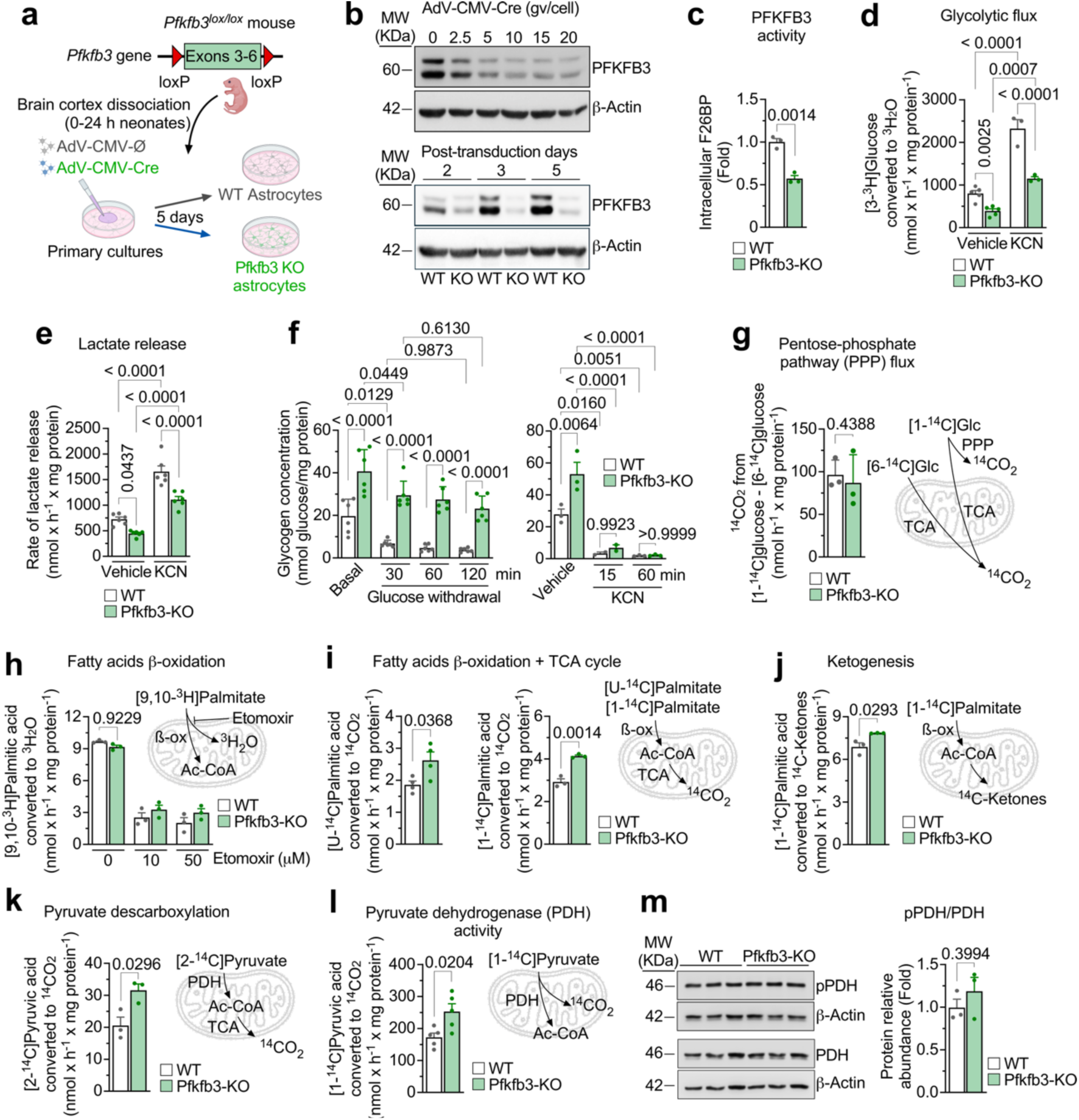
Inhibited glycolysis in Pfkfb3-KO astrocytes boosts pyruvate decarboxylation and TCA cycle activity. **(a)** Strategy used to obtain *Pfkfb3* knockout astrocytes in primary culture. Created with BioRender.com. **(b)** *Upper panel*: representative Western blot of primary astrocytes showing PFKFB3 protein levels across a gradient of AdV-CMV-Cre viral genomes per cell (gv/cell); β-Actin was used as loading control. *Lower panel*: Western blot against PFKFB3 in WT and Pfkfb3-KO astrocytes at indicated time points (days) post-transduction (15 gv/cell). In all next experiments, 15 gv/cell for 5 days was used. (See also Extended Data Fig. 2a). **(c)** Fructose-2,6-bisphosphate (F26BP) levels, a proxy for PFKFB3 activity, in WT and Pfkfb3-KO primary astrocytes. Data are mean ± S.E.M. *P* value is indicated; n=3 biologically independent samples per genotype; Unpaired Student’s t-test, two-tailed. **(d)** Glycolytic flux in WT and Pfkfb3-KO primary astrocytes untreated (vehicle) or treated with potassium cyanide (KCN, 1 mM). Data are mean ± S.E.M. *P* values are indicated; n=5 (vehicle) or 3 (KCN) biologically independent samples per genotype; two-way ANOVA followed by Tukey. (See also Extended Data Fig. 2b). **(e)** Lactate release from WT and Pfkfb3-KO primary astrocytes untreated (vehicle) or treated with KCN. Data are mean ± S.E.M. *P* values are indicated; n=6 biologically independent samples per genotype; two-way ANOVA followed by Tukey. **(f)** Glycogen concentrations in WT and Pfkfb3-KO primary astrocytes under basal conditions (with glucose) or following glucose withdrawal at the indicated times (*left*). *Right*: glycogen concentrations in WT and Pfkfb3-KO primary astrocytes untreated (vehicle) or treated with KCN for the indicated times. Data are mean ± S.E.M. *P* values are indicated; n=6 (*left panel*), 2 (*right panel*, KCN 15 min) or 3 (*right panel*, vehicle and KCN 60 min) biologically independent samples per genotype; two-way ANOVA followed by Tukey. **(g)** Pentose-phosphate pathway (PPP) flux in WT and Pfkfb3-KO primary astrocytes. Data are mean ± S.E.M. *P* value is indicated; n=3 biologically independent samples per genotype; Unpaired Student’s t-test, two-tailed. TCA, tricarboxylic acid cycle. (See also Extended Data Fig. 2b). **(h)** Fatty acid β-oxidation in WT and Pfkfb3-KO primary astrocytes treated with increasing concentrations of etomoxir. Data are mean ± S.E.M. *P* value is indicated; n=3 biologically independent samples per genotype; two-way ANOVA followed by Tukey. **(i)** Combined fatty acid ß-oxidation and TCA cycle flux in WT and Pfkfb3-KO primary astrocytes using either [U-^14^C]- (*left*) or [1-^14^C]palmitic acid (*right*). Data are mean ± S.E.M. *P* values are indicated; n=4 (U-^14^C) or 3 (1-^14^C) biologically independent samples per genotype; Unpaired Student’s t-test, two-tailed. **(j)** Ketogenesis rate in WT and Pfkfb3-KO primary astrocytes. Data are mean ± S.E.M. *P* value is indicated; n=3 biologically independent samples per genotype; Unpaired Student’s t-test, two-tailed. **(k)** Pyruvate decarboxylation at the TCA cycle in WT and Pfkfb3-KO primary astrocytes. Data are mean ± S.E.M. *P* value is indicated; n=3 biologically independent samples per genotype; Unpaired Student’s t-test, two-tailed. **(l)** Pyruvate decarboxylation to acetyl-CoA at pyruvate dehydrogenase (PDH activity) in WT and Pfkfb3-KO primary astrocytes. Data are mean ± S.E.M. *P* value is indicated; n=5 biologically independent samples per genotype; Unpaired Student’s t-test, two-tailed. **(m)** Western blot against phosphorylated PDH (pPDH) and total PDH protein in WT and Pfkfb3-KO primary astrocytes; β-Actin was used as loading control (*left*). Quantification of the pPDH/PDH ratio (*right*). Data are mean ± S.E.M. *P* value is indicated; n=3 biologically independent samples per genotype; Unpaired Student’s t-test, two-tailed.

Given that astrocytic fatty acid ß-oxidation may take over the energetic -and consequently survival-function(s) of glycolysis^26^, we sought to investigate ß-oxidation of fatty acids upon *Pfkfb3* loss. As shown in **Fig. 2h**, the mitochondrial production of ^3^H_2_O from [9,10-^3^H]palmitate, a *bona fide* measure of acyl-coenzyme A (acyl-CoA) to acetyl-CoA conversion -i.e., strictly fatty acid ß-oxidation^19^-, was unaltered in *Pfkfb3* KO astrocytes. Intriguingly, the rates of [U-^14^C]- and [1-^14^C]palmitate oxidation to ^14^CO_2_, which measures fatty acid ß-oxidation *plus* acetyl-CoA decarboxylation in the TCA cycle, increased by ∼1.5 fold in the glycolytically-impaired *Pfkfb3* KO astrocytes (**Fig. 2i**); however, ketogenesis, i.e., the conversion of fatty acids-derived acetyl-CoA into ketone bodies, as determined by the rate of incorporation of [1-^14^C]palmitate into [^14^C]ketones^19,27^, was modestly increased (∼1.1-fold) (**Fig. 2j**). Together, these data indicate that, upon glycolysis reduction in *Pfkfb3* KO astrocytes, fatty acids catabolism to acetyl-CoA (ß-oxidation) remains unaltered, but acetyl-CoA decarboxylation *via* the TCA cycle is up-regulated. Whilst these data apparently contrast with those of *Drosophila*, in which ß-oxidation takes over the metabolic functions of a complete abolishment of glycolysis^26^, glycolysis is only partially reduced in *Pfkfb3* KO astrocytes, suggesting that a proportion of pyruvate might be preserved for mitochondrial utilization. To test this possibility, we investigated mitochondrial pyruvate catabolism. As shown in **Fig. 2k**, the rate of [2-^14^C]pyruvate decarboxylation, a process that takes place at α-ketoglutarate dehydrogenase complex hence indicating acetyl-CoA decarboxylation at the TCA cycle, increased in astrocytes lacking *Pfkfb3*. Notably, the flux through pyruvate dehydrogenase (PDH) complex increased at the same extent, as revealed by [1-^14^C]pyruvate decarboxylation (**Fig. 2l**), indicating that decreased glycolysis in *Pfkfb3* KO astrocytes leads to enhanced mitochondrial pyruvate oxidation through PDH complex followed by TCA cycle. Wild type astrocytic PDH was highly phosphorylated, and this was unchanged in *Pfkfb3* KO astrocytes (**Fig. 2m**), indicating that the increased flux through PDH complex likely results from the law of mass action, rather than by a post-translational mechanism. This is coherent with the observed decrease in lactate release by primary astrocytes (**Fig. 2e**), decreased pyruvate and lactyl-valine, and increased α-ketoglutarate (**Fig. 1h**) in the brain. Altogether, these data show that *Pfkfb3* KO astrocytes undergo a metabolic rewiring consisting of boosted mitochondrial pyruvate decarboxylation leading to reduced lactate formation.

PDH and TCA cycle activities reduce NAD^+^ to NADH(H^+^), but our data (**Extended Data Fig. 3a**) show that NAD^+^/NADH(H^+^) ratio remains unchanged in *Pfkfb3* KO astrocytes, suggesting the concomitant activation of a mitochondrial NADH(H^+^) oxidizing machinery. Since the electron transport chain (ETC) activity, at the level of complex I, largely accounts for mitochondrial NADH(H^+^) oxidation, we then assessed the ETC activity as the oxygen consumption rate (OCR). Our data revealed a specific enhancement in mitochondrial respiration, as judged by the increased basal and maximal OCR, spare respiratory capacity and ATP-linked OCR with a negligible -although statistically significant-increase in non-mitochondrial OCR in *Pfkfb3* KO astrocytes (**Fig. 3a**). Furthermore, blue-native gel electrophoresis of mitochondrial extracts revealed that complex I super assembled with complex III in *Pfkfb3* KO astrocytes (**Fig. 3b**), a conformation known to improve electron flux through complex I in the ETC^28^. In contrast, as expected^28^, in *Pfkfb3* KO astrocytes complex II-mediated mitochondrial respiration decreased (**Fig. 3c**), as it did the production of mitochondrial ROS (**Extended Data Fig. 3b**) -which are mostly produced by non-super assembled complex I^19,29^. Altogether, these data imply that enhanced PDH and TCA cycle activities in glycolysis-deficient *Pfkfb3* KO astrocytes is coupled to increased complex I-driven ETC stimulation, resulting in NADH(H^+^)/NAD^+^ ratio preservation.

**Fig. 3.**
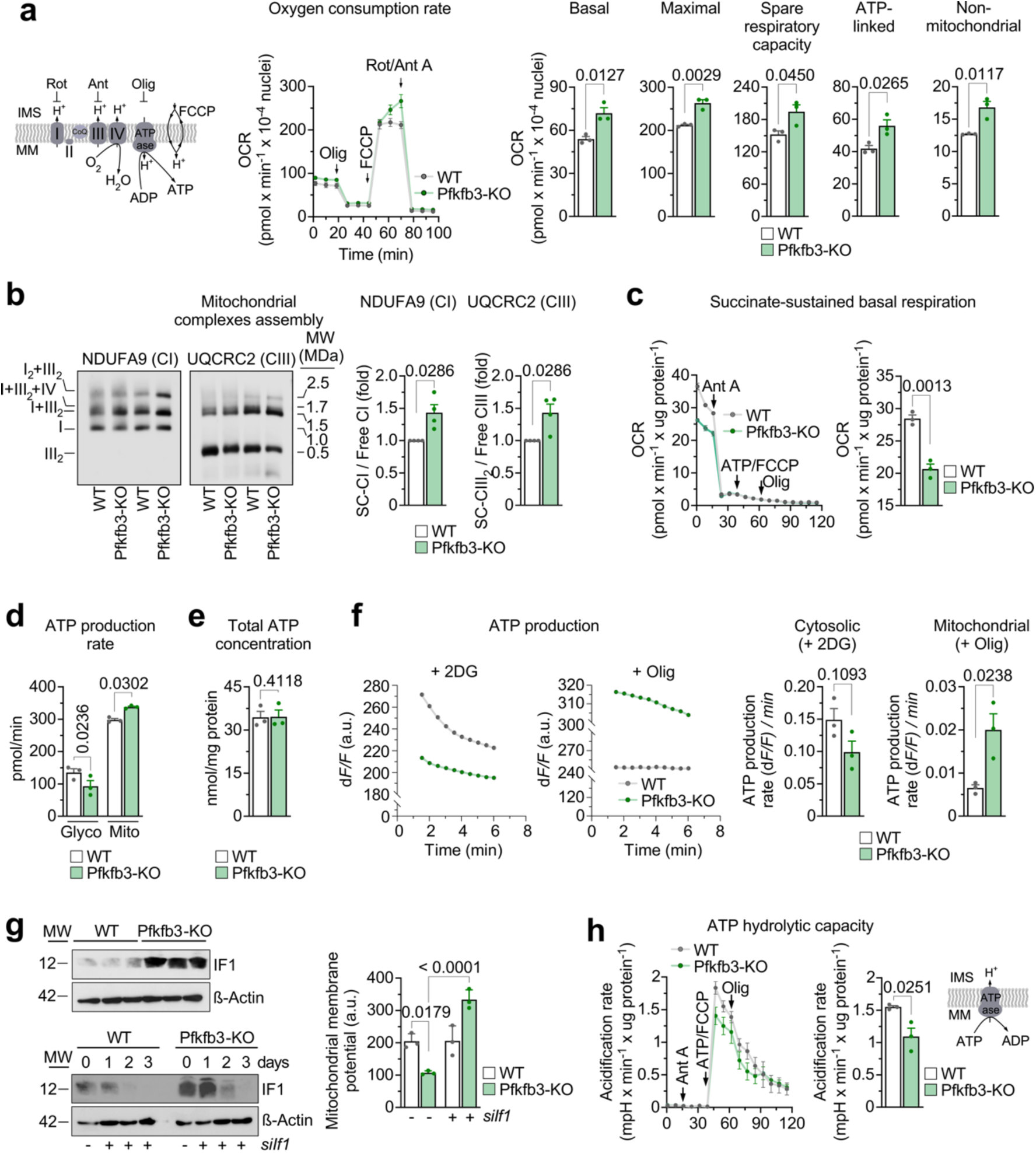
Impaired glycolysis in Pfkfb3-KO astrocytes switches mitochondria from hydrolysis to synthesis of ATP and promotes respiration. **(a)** Oxygen consumption rate (OCR) analysis in WT and Pfkfb3-KO astrocytes in primary culture. *Left*: schematic representation indicating the inhibitors sites of action (Rot, rotenone to inhibit complex I; Ant, antimycin to inhibit complex III; Olig, oligomycin to inhibit ATP synthase; FCCP, uncoupler); *middle*: OCR traces; *right*: parameters quantification. Data are mean ± S.E.M. *P* values are indicated; n=3 biologically independent samples per genotype; Unpaired Student’s t-test, two-tailed. (See also Extended Data Fig. 3a). (IMS, intermembrane space; MM, mitochondrial matrix). **(b)** Analysis of the mitochondrial respiratory chain complexes assembly. *Left*: Blue-native gel electrophoresis images showing complex I (CI)- or complex III (CIII) containing supercomplexes (SC-CI, SC-CIII) and free CI and CIII in WT and Pfkfb3-KO astrocytes. *Right*: Quantification of SC *versus* free CI and CIII, normalized to WT. Data are mean ± S.E.M. *P* values are indicated; n=4 biologically independent samples per genotype; Unpaired Mann-Whitney test, two-tailed. **(c)** Succinate-driven basal respiration in disrupted mitochondrial preparations from WT and Pfkfb3-KO astrocytes. *Left*: OCR trace. *Right*: OCR quantification. Data are mean ± S.E.M. *P* value is indicated; n=3 biologically independent samples per genotype; Unpaired Student’s t-test, two-tailed. **(d)** ATP production rates from glycolysis (*Glyco*) and mitochondria (*Mito*) in WT and Pfkfb3-KO astrocytes. Data are mean ± S.E.M. *P* values are indicated; n=3 biologically independent samples per genotype; Unpaired Multiple t-test. **(e)** Total cellular ATP content in WT and Pfkfb3-KO astrocytes. Data are mean ± S.E.M. *P* value is indicated; n=3 biologically independent samples per genotype; Unpaired Student’s t-test, two-tailed. (See also Extended Data Fig. 3a). **(f)** ATP production dynamics measured as fluorescence change (dF/F) over time upon inhibition of glucose utilization (+2DG, 2-deoxyglucose) or mitochondrial ATP synthase (+Olig, oligomycin) (*left*). *Right*: quantifications of the ATP production rate by cytosolic (glycolysis; + 2DG) or mitochondrial (ATP synthase; + Olig) pathways, in WT and Pfkfb3-KO astrocytes. Data are mean ± S.E.M. *P* values are indicated; n=3 biologically independent samples per genotype; Unpaired Student’s t-test, two-tailed. **(g)** *Upper panel*: Western blot against mitochondrial ATP synthase inhibitory factor 1 (IF1) in WT and Pfkfb3-KO astrocytes; β-Actin was used as loading control. *Lower panel*: Western blot against IF1 in WT and Pfkfb3-KO astrocytes transfected with small interfering RNA against If1 (siIf1). *Right panel*: mitochondrial membrane potential in WT or Pfkfb3-KO astrocytes treated with scrambled siRNA (- *siIf1*) or with (+) *siIf1*. Data are mean ± S.E.M. *P* values are indicated; n=3 biologically independent samples per genotype; two-way ANOVA. **(h)** ATP hydrolytic capacity as assessed by the kinetics of medium acidification in WT and Pfkfb3-KO astrocytes. *Left*: traces. *Right*: quantification of the acidification rates. The scheme inset illustrates coupling of ATP hydrolysis to proton pumping. Data are mean ± S.E.M. *P* value is indicated; n=3 biologically independent samples per genotype; Unpaired Student’s t-test, two-tailed.

Given that ETC activity is coupled to ATP production, we then investigated the impact of enhanced ETC activity in *Pfkfb3* KO astrocytes on cellular ATP synthesis. We found that glycolysis-deficient *Pfkfb3* KO astrocytes decreased glycolytic-but increased mitochondrial ATP production rate (**Fig. 3d**) without altering total ATP concentrations (**Fig. 3e**), which was confirmed by real-time fluorescence imaging (**Fig. 3f**). The protein abundance of ATP synthase inhibitor factor-1 (IF1), an inhibitor of the reverse mode ATP synthase activity -ATP hydrolytic-, was found dramatically increased in *Pfkfb3* KO astrocytes (**Fig. 3g**). More interestingly, IF1 knockdown largely overcame the decreased mitochondrial membrane potential (Δψ_m_) caused by *Pfkfb3* loss (**Fig. 3g**). Thus, upon *Pfkfb3* loss there is a subsequent decrease in glycolytic ATP production, which does not meet the mitochondrial needs to sustain the Δψ_m_ that otherwise can be maintained in the absence of IF1. Given that these data suggest that Δψ_m_ is sustained in *Pfkfb3* KO astrocytes by the ETC, we next determined the reverse-mode ATP synthase activity. Consistent with the increased IF1 abundance (**Fig. 3g**), *Pfkfb3* loss decreased the ATP hydrolytic activity (**Fig. 3h**), confirming that ATP synthase activity is switched from reverse to forward mode. Altogether, astrocytes with weakened glycolysis sustain cellular ATP by enhancing mitochondrial pyruvate oxidation through PDH and TCA cycle, thus fueling the ETC to maintain Δψ_m_ and feed the forward-mode of ATP synthase.

Next, we aimed to elucidate whether this metabolic reprogramming accounts for the cognitive impairment observed in the astrocytic-specific *Pfkfb3* KO mice. To do so, we knocked down the catalytic, pyruvate dehydrogenase-1a subunit (*Pdh1a*) of the PDH complex. This treatment was efficient, as judged by the reversal of the increase in [2-^14^C]pyruvate decarboxylation (**Fig. 4a**), the decrease in lactate release (**Fig. 4b**) and the enhanced mitochondrial respiration (**Fig. 4c**) observed in *Pfkfb3* KO astrocytes. Of note, *in vivo* knockdown of *Pdh1a* specifically in astrocytes (**Fig. 4d**, *left***; Extended Data Fig. 4a,b**) prevented the short-term memory loss observed in astrocyte-specific *Pfkfb3* KO mice (**Fig. 4d**, *middle and right panels*), as judged by the novel object recognition test. This result indicates that an enhancement in astrocytic mitochondrial pyruvate oxidation accounts for memory impairment in astrocytic-specific *Pfkfb3* KO mice. In addition, since astrocytic *Pfkfb3* loss causes a decrease in the release of lactate (**Figs. 2e**, **4b**), we sought to investigate whether the decreased astrocytic-derived lactate observed in astrocytic-specific *Pfkfb3* KO mice could be a key contributing factor in the loss of memory. To address this issue, astrocytic-specific *Pfkfb3* KO mice received a single systemic dose of lactate, according to a previously validated protocol^30^, and cognition was then assessed. Whilst lactate administration had no effect on short-term memory in wild type mice (**Extended Data Fig. 4c**), it fully rescued the memory loss observed in the astrocytic-specific *Pfkfb3* KO mice (**Fig. 4e**, *middle and right panels*). To ascertain whether this protective effect was due to lactate uptake by neurons, we also administered lactate to astrocytic-specific *Pfkfb3* KO mice in which monocarboxylate transporter-2 (*Mct2*) -the predominant lactate transporter in neurons^31^-was previously knocked down specifically in neurons (**Fig. 4e***, left panel***; Extended Data Fig. 4d**). The results show that neuron-specific *Mct2* knockdown abolished the protective effect of lactate administration on cognition in astrocyte-specific *Pfkfb3* KO mice (**Fig. 4e**, *middle and right panels*), but not in wild type animals (**Extended Data Fig. 4c**). To confirm that the behavioral outputs observed upon these treatments correlate with neuronal function, we assessed *cFos* and *Arc* mRNA expression in the hippocampus. The results confirmed decreased *cFos* and *Arc* expression -indicating neuronal dysfunction-in astrocytic-specific *Pfkfb3* KO mice, which was abolished by either *Pdh1a* knockdown or lactate administration -unless *Mct2* was knocked down (**Extended Data Fig. 4e**).

**Fig. 4.**
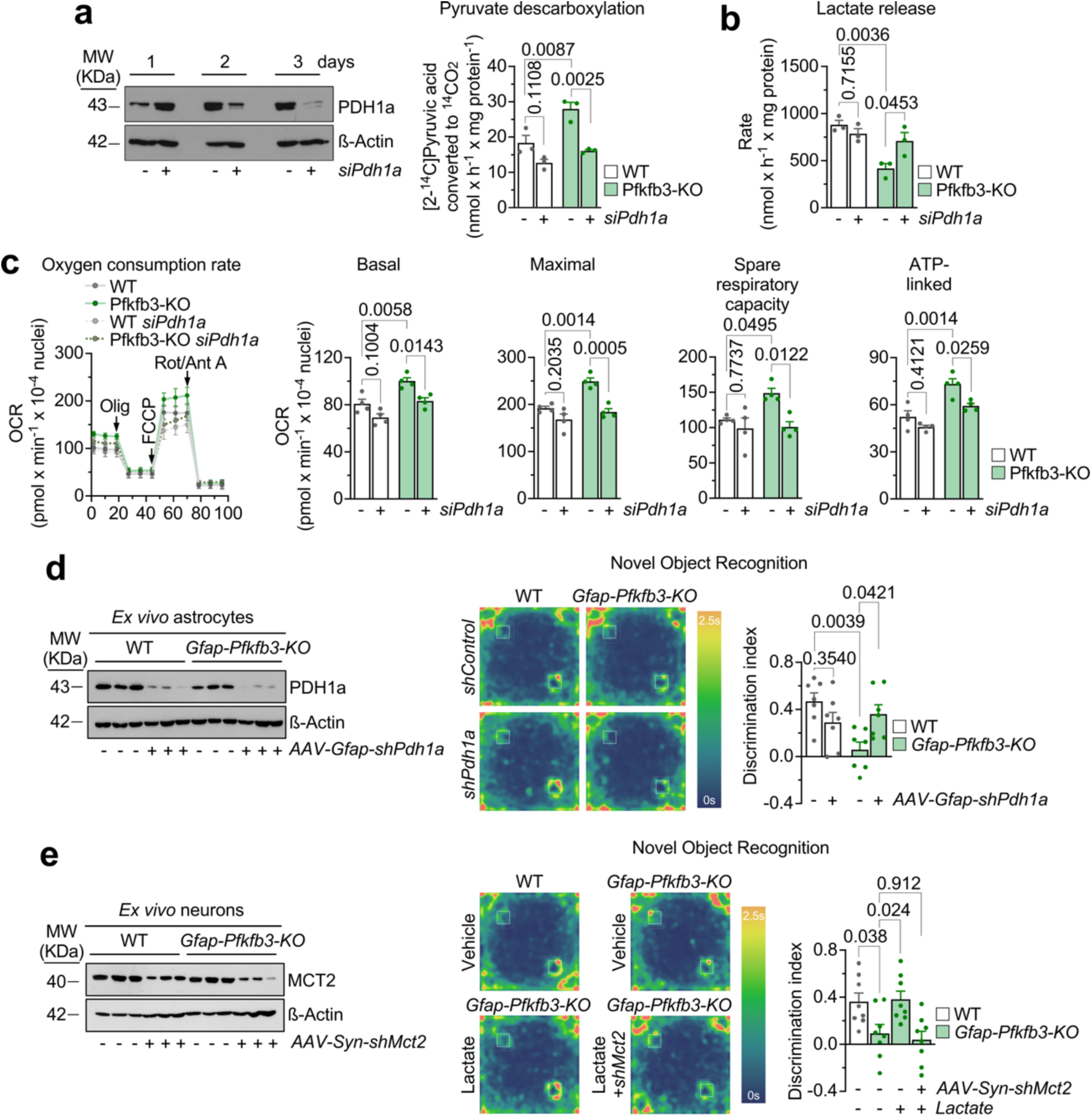
Astrocyte-specific PDH silencing rescues cognitive deficits of *Gfap-Pfkfb3-KO* mice *via* lactate transport to neurons. **(a)** Western blotting against PDH1a in primary astrocytes transfected with *siPdh1a* in primary astrocytes at different time points; ß-actin was used a loading control (*left*). *Right*: pyruvate decarboxylation rate in WT and Pfkfb3-KO astrocytes transfected with scrambled (-) or *siPdh1a*. Data are mean ± S.E.M. *P* values are indicated; n=3 biologically independent samples per genotype and condition; two-way ANOVA followed by Tukey. **(b)** Rate of lactate release by WT and Pfkfb3-KO astrocytes transfected with scrambled (-) or *siPdh1a*. Data are mean ± S.E.M. *P* values are indicated; n=3 biologically independent samples per genotype and condition; two-way ANOVA followed by Tukey. **(c)** OCR in WT and Pfkfb3-KO astrocytes transfected with scrambled (-) or *siPdh1a*. Data are mean ± S.E.M. *P* values are indicated; n=4 biologically independent samples per genotype and condition; two-way ANOVA followed by Tukey. **(d)** Western blotting against PDH1a in astrocytes immunomagnetically isolated from WT or *Gfap-Pfkfb3-KO* mice 10 days after *in vivo* transduction with *AAV-Gfap-shPdh1a* (*left*). Novel object recognition test average occupancy heat maps in WT and *Gfap-Pfkfb3-KO* mice treated or not with *AAV-Gfap-shPdh1a* (*middle*). Discrimination index quantification (*right*). Data are mean ± S.E.M. *P* values are indicated; n=7 mice per genotype and treatment; two-way ANOVA followed by Tukey. (See also Extended Data Fig. 4a,b). **(e)** Western blotting against MCT2 in astrocytes immunomagnetically isolated from WT or *Gfap-Pfkfb3-KO* mice 10 days after *in vivo* transduction with *AAV-Gfap-shMct2* (*left*). Novel object recognition test representative occupancy heat maps in WT and *Gfap-Pfkfb3-KO* mice treated or not with *AAV-Gfap-shMct2* (*middle*). Discrimination index quantification (*right*). Data are mean ± S.E.M. *P* values are indicated; n=8 mice per genotype and treatment; one-way ANOVA followed by Dunnett. (See also Extended Data Fig. 4c,d,e).

In conclusion, here we show that astrocytes require a high glycolytic phenotype, sustained by Pfkfb3 activity, to maintain cognition through a mechanism engaging glycolytically derived ATP with mitochondrial consumption *via* the reverse mode of ATP synthase. By hydrolyzing this ATP, astrocytic mitochondria maintain Δψ_m_ thus preventing mitochondria from using pyruvate to sustain TCA cycle and ETC activities (**Extended Data Fig. 4f**). Along with glycolysis, Pfkfb3 bolsters glycogen catabolism, which contributes to the prevalent release of lactate by astrocytes^23,24^ that takes place at the expense of reduced mitochondrial pyruvate uptake. Notably, although through a different mechanism, fatty acid ß-oxidation^19^ also contributes to cognition by modulating mitochondrial competence. Thus, ß-oxidation in astrocytes is primarily ketogenic and directly fuels ubiquinone through the electron transfer flavoprotein, which reduces the formation of supercomplexes between complexes I and III^19^, favoring their isolated versions thus affecting complex I rate at oxidizing NADH(^+^)^32^ and inducing higher ROS production^29^. However, according to the present data, Pfkfb3-promoted glycolysis facilitates pyruvate to lactate conversion, weakening in this way TCA cycle, which in turn attenuates complex I contribution to NADH(H^+^) oxidation and ROS formation. Therefore, both metabolic pathways, namely Pfkfb3-promoted glycolysis and fatty acid ß-oxidation^19^ converge in a unifying mechanism that may cooperatively work to attenuate mitochondrial pyruvate utilization. This metabolic collaboration would ensure the supply of readily oxidizable metabolites, such as glycolysis-derived lactate^5,9,30^ and ß-oxidation-derived ketone bodies^27,33^ to neighbor, glycolytically-weak^1,34,35^ neurons to fuel their energy needs. In good agreement with this notion, here we show that the memory loss observed in astrocytic-specific *Pfkfb3*-KO mice can be fully rescued by concomitant silencing of *Pdh1a* selectively in astrocytes and by lactate administration -unless the neuron-specific lactate transporter, *Mct2*, was silenced selectively in neurons of the astrocyte-specific *Pfkfb3*-KO mice. Furthermore, our work provides a specific molecular mechanism that resolves *in vivo* the long-lasting debate on the occurrence of an astrocyte-to-neuron lactate shuttle^5,9,30^. Thus, our data provide an explanation as to why neurons, which display a weak glycolytic activity^1,2,35^, must rely on neighbor astrocytes for metabolic support and hence preventing neurological diseases such as multiple sclerosis, ischemia, and cognitive dysfunction^8,35–41^. Our findings identify astrocytic Pfkfb3 as a potentially valuable metabolic target for therapeutic intervention in neurological disorders.

## Methods

### Pfkfb3^lox/lox^ mice

All protocols were performed according to the European Union Directive 86/609/EEC and Recommendation 2007/526/EC, regarding the protection of animals used for experimental and other scientific purposes, enforced in Spanish legislation under the law 6/2013. Protocols were approved by the Bioethics Committee of the University of Salamanca in accordance with the Spanish legislation (RD53/2013). All mice used in this study were of the C57Bl/6J background. *Pfkfb3^lox/lox^*mice was generated by introducing two loxP sites flanking a segment comprising exon 3 and 6 of *Pfkfb3* gene *via* homologous recombination in embryonic stem cells under a C57BL/6J background^18^. Animals were bred at the Animal Experimentation Facility of the University of Salamanca in cages (maximum of five animals per cage) with a 12 h light-dark cycle (light from 08:00 h). The humidity was 45–65%, and the temperature was 20-25 °C. Animals were fed *ad libitum* with a standard solid diet (Envigo-Harlan Tekland Global 18% Protein Rodent Diet, USA; 18% proteins, 3% lipids, 58.7% carbohydrates, 4.3% cellulose, 4% minerals, and 12% humidity) and free access to water.

### *In vivo* generation of astrocyte-specific *Pfkfb3* knockout mice

This was carried out using a validated adeno-associated virus (AAV) strategy^20^. Essentially, AAV particles of the PHP.eB capsid (serotype), known to efficiently transduce the central nervous system *via* intravenous injection^42^, expressing Cre recombinase driven by the astrocyte-specific short glial-fibrillary acidic protein (GFAP) promoter (PHP.eB-AAV-gfaABC_1_D-Cre-GFP) were administered intravenously (50 µl aliquots of a phosphate-buffered saline solution containing 0.001% Pluronic® F-68, Sigma-Aldrich, and 1 × 10^11^ viral genomes, VG) through the retro-orbital sinus to 3 months-old *Pfkfb3^lox/lox^* male mice under a brief sevoflurane anesthesia (Sevorane, AbbVie, Spain, at 6% for initiation followed by ∼3% for maintenance in air with supplement O_2_ and NO_2_ -0.4 and 0.8 litres/min, respectively-using a gas distribution column, *Hersill H-3*, Spain, and a vaporizer, *InterMed Penlons* Sigma Delta, UK). We used the retro-orbital sinus intravenous route because of the higher success rate observed when compared with the tail or temporal ones^43^. Siblings of wild-type (WT) mice received equivalent amounts of the same AAV particles that did not harbor Cre recombinase. Mice were used from 6 weeks after AAV injections. The efficacy of this approach at promoting recombination exclusively in astrocytes was previously confirmed in our laboratory using *Cpt1a^lox/lox^* mice^19^.

### Virus generation

Adeno-associated vectors of the PHP.eB serotype containing the sequence of a shRNA directed against the *Slc16a7* gene, which codes for the Mct2 protein, or directed against the *Pdha1* gene, were produced by Vector Builder Inc. The miR30-based shRNA system was used to subject the shRNA expression to the neuronal-specific Syn promoter or the astrocytic GFAP promoter, respectively. The shRNA sequence used to knock-down *Mct2* and *Pdha1* were: *shSlc16a7*: 5’-CTAGGATTAATAGCCAACACTATAGTGA-AGCCACAGATGTATAGTGTTGGCTATTAATCCTAT-3’; *shPdha1*: 5’-ACCAGA-CTTACCGCTACCATGGTAGTGAAGCCACAGATGTACCATGGTAGCGGTAAG TCTGGG-3’.

### Treatment of mice with sodium lactate

Mice received a single intraperitoneal injection of sodium lactate (L7022, Sigma-Aldrich; 534 mmol/L, 4 μL/g mice) dissolved in NaCl 0.9 % as the vehicle 20 min before the behavioral testing to ensure maximum brain lactate concentration. Control mice received an identical volume of the vehicle^44–46^.

### Primary cultures of astrocytes

Astrocytes in primary culture were obtained from the cortex of 0-24 h old *Pfkfb3^lox/lox^* mouse^18^ neonates^47^. Cell suspension was seeded in 175 cm^2^ plastic flasks in low glucose (5.5 mM) Dulbecco’s Modified Eagle’s Medium (DMEM) supplemented with 10% fetal bovine serum and 4 mM glutamine and incubated at 37 °C in a humidified 5% CO_2_-containing atmosphere. To detach non-astrocytic cells, after 7 days *in vitro* (DIV), the flasks were shaken at 150 r.p.m. overnight. The supernatant was discarded, and the attached, astrocyte-enriched cells were reseeded at 0.8-1 × 10^5^ cells/cm^2^ in the appropriate plates. Cells were used at 14 DIV. Immunocytochemistry against a neuronal (β-Tubulin III: 1/300; T2200; Sigma), astrocytic (GFAP: 1/800; AB5541; Millipore), oligodendrocytes (O4; 1/300; from mouse hybridoma kindly donated by Isabel Fariñas’ laboratory) and microglial marker (CD45; 1/200; 553076; BD) was performed in order to determine the purity of the cultures, which was ∼100% astrocytes^19^.

### Generation of *Pfkfb3* knockout astrocytes in primary culture

This was carried out by transducing 9 DIV primary astrocytes, obtained from *Pfkfb3^lox/lox^* mice^18^, with adenoviral particles harboring Cre recombinase driven by the ubiquitous cytomegalovirus (CMV) promoter (AdV-CMV-Cre). The were purchased from the viral repository of the University of Iowa (VVC-U of Iowa-5 for AdV-CMV-Cre; VVC-U of Iowa-272 for AdV-CMV-Ø). Astrocytes from the same cultures transduced with equivalent amounts of the same AdV lacking Cre recombinase (AdV-CMV-Ø) were used as WT astrocytes. Cells were used 5 days after transduction. Potassium cyanide (KCN) was used in some experiments as complex IV of the ETC, leading to energy depletion and an increase in the glycolytic pathway. KCN was weighed and diluted extemporaneously in the incubation medium itself; the times are indicated in each case. A concentration of 1 mM was regularly used to produce this increase in glycolysis without completely inhibiting respiration.

### Genotyping and recombination event by polymerase chain reaction (PCR)

For *Pfkfb3^lox/lox^* genotyping, a PCR with the following primers was performed; primer 1, 5’-CGAGACAATGTTCCATAGCTTGAATG -3’ (forward) and primer 2, 5’-CAGGCCCAGACCAAGGACAGC-3’ (reverse), resulting in a 500 bp band for *Pfkfb3^lox/lox^* mice and 400 bp for wild type^18^. To enable distinguishing the non-excised recombined allele from the Cre-mediated excised allele, a PCR with the following primers was performed: primer 1, 5’-CGAGACAATGTTCCATAGCTTGAATG -3’ (forward) and primer 3, 5’-GGCTGTGCTATAGTGTGAGAATCCT-3’ (reverse). PCR conditions were 5 min at 94°C, 35 cycles of 30 s at 94°C, 30 s at 55°C, 30 s at 72°C, and final extension of 10 min at 72°C. Products were resolved in 3% agarose gel using the 100 bp and 1 Kb DNA ladder plus (Thermo Fisher Scientific).

### Small interference RNA

To obtain specific protein knockdown, we used the following small interference RNA (siRNAs; only the forward strand shown), ATPase inhibitory factor 1 (Atpif1) 5’AGAUGAGAUUGACCACCAUtt-3’(Mus musculus Gen ID:11983); Pyruvate dehydrogenase (lipoamide) subunit alpha 1 (*Pdha1*), 5’GACUUACCGCUACCAUGGAtt-3’ (Mus musculus Gen ID:18597), and siRNA control (siControl; 4390843; Thermo Fisher). Transfections with siRNAs were performed with *Lipofectamine RNAiMAX reagent* (Thermo Fisher) according to the manufactureŕs protocol using a siRNA final concentration of 9 nM.

### Immunomagnetic purification of astrocytes and neurons from adult brain

Mouse adult brain (minus cerebellum and olfactory bulb) was dissociated using the adult mouse brain dissociation kit (Miltenyi Biotec). The tissue, once clean, was fragmented with a sterile scalpel in 2 ml per hemisphere of a disintegration solution (Earle’s Balanced Salt Solution, EBSS, 116 mM NaCl, 5.4 mM KCl, 1.5 mM MgSO_4_, NaHCO_3_ 26 mM, NaH_2_PO_4_•2H_2_O 1.01 mM, glucose 4 mM, phenol red 10 mg/l, supplemented with albumin 14.4 μl/ml and DNase type I 26 μl/ml, pH 7.2, trypsin 10.8μl/ml), and it was trypsinized at 37°C in a thermostated bath for 5 minutes, shaking frequently to avoid decantation of the tissue. It was further mechanically disintegrated by trituration using a 5 ml serological pipette for 5 times. Then, the suspension was returned to the thermostated bath for 10 minutes, shaking frequently. Trypsin activity was stopped by adding 10% fetal serum, before centrifuging the tissue at 700 g for 5 minutes in a microfuge at 4°C. Once the enzymatically disintegrated tissue had been decanted, the pellet was resuspended in a trypsin-free disintegration solution (EBSS + 13 μl/ml DNase + 20 μl/ml albumin) for mechanical trituration using a Pasteur pipette. Approximately 5 passages were performed per volume of 4 ml and per hemisphere. The supernatant was centrifuged for 3 min at 700 g and the number of cells in the pellet was counted. Once a homogeneous suspension of individualized adult neural cells was achieved, cell population separations were performed using MACS® Technology using either the astrocyte-specific anti-ACSA-2 Microbead Kit or the neuron-specific Neuron Isolation Kit, according to manufacturer’s protocol (MACS® technology). We confirmed the identity of the isolated fractions by Western blotting against astrocytic (GFAP), neuronal (MAP2)-specific markers, and the purity with microglial (Iba1) and oligodendroglial (OLIG2)-specific markers^19^.

### Fructose-2,6 bisphosphate (F26BP) determinations

For F26BP determinations, cells were lysed in 0.25 N NaOH and centrifuged (20,000 x *g*, 20 min). An aliquot of the homogenate was used for protein determination, and the remaining sample was heated at 80 °C (5 min), centrifuged (20,000 x *g*, 10 min, 4°C), and the resulting supernatant used for the enzymatic determination of F26BP concentrations using F26BP standards, as previously described^48^. The F26BP standard was prepared by incubating a 20 mM solution of D-fructose-1,2-cyclic 6-bisphosphate (Sigma; catalogue number 68872) in 0.5 M NaOH at 37 °C for 30 min. To determine F26BP levels, 20 µl of the supernatant was diluted 1:20 in 0.05 M HCl. In parallel, a non-acidified reaction was carried out by diluting the supernatant 1:10 in 0.05 M NaCl. After a 10 min incubation at room temperature, which quantitatively converts F26BP into fructose-6-phosphate (F6P), the samples were neutralized with 0.1 M NaOH and used to determine F6P in a buffer containing 25 mM HEPES (pH 8.2), 25 mM KCl, 2.5 mM magnesium acetate, 1 mM dithiothreitol, and 0.25 mM NADP^+^ in the presence of glucose-6-phosphate dehydrogenase and phosphoglucose isomerase. F26BP relative levels in the samples were assessed by the coupled enzymatic activities of 6-phosphofructo-1-kinase (Pfk1) (Sigma; catalogue number F2258) in the presence of 1 mM fructose-6-phosphate (Sigma; catalogue number F3627) and 0.5 mM pyrophosphate (Sigma; catalogue number P8010), aldolase (Sigma; catalogue number A8811), and triose-phosphate isomerase/glycerol-3-phosphate dehydrogenase (Sigma; catalogue number G1881). This reaction generates glycerol-3-phosphate and oxidizes NADH (Sigma; catalogue number N8129), producing a reduction in the absorbance at 340 nm that is monitored spectrophotometrically. F26BP relative concentrations are calculated in the samples by comparing the fold-activation that they cause on Pfk1 activity.

### Determination of metabolic fluxes

To assess glucose, fatty acid, and pyruvate oxidative fluxes, we used radiometric approaches. To do this, astrocytes were seeded in 8 cm^2^ flasks (353108, Fisher) hanging a microcentrifuge tube containing either 1 ml benzethonium hydroxide (Sigma) (for ^14^CO_2_ equilibration) or 1 ml H_2_O (for ^3^H_2_O equilibration). To assess glycolytic flux from adult brain cell suspensions were placed in 25-cm^2^ glass flasks harboring a central well with a tube containing 1 ml H_2_O. All incubations were carried out in DMEM (D5030, 8.3 g/L, pH 7.4, Sima) supplemented with glutamine 4 mM, pyruvate 1 mM, glucose 5.5 mM, and HEPES 5 mM at 37 °C in the air-thermostatized chamber of an orbital shaker. The glycolytic flux was measured by assaying the rate of ^3^H_2_O production from [3-^3^H]glucose using 2 μCi/ml of D-[3-^3^H]glucose in DMEM for 3 h ^1^. The rate of fatty acids ß-oxidation -i.e., the conversion of fatty acids to acetyl-Coenzyme A-was performed by analyzing the rate of ^3^H_2_O production from [9,10-^3^H]palmitate (1 μCi/ml) in DMEM supplemented with 10 μM palmitate for 3 h^49–51^. Both the glycolytic flux and ß-oxidation, after incubations were terminated with 0.2 ml 20 % perchloric acid, the cells were further incubated for 72 h to allow for ^3^H_2_O equilibration with H_2_O present in the central microcentrifuge tube. The ^3^H_2_O was then measured by liquid scintillation counting *Tri-Carb 4810 TR* (PerkinElmer). To measure the carbon flux from fatty acids to CO_2_, -which jointly assesses ß-oxidation and acetyl-CoA decarboxylation in the TCA cycle-, cells were incubated in DMEM supplemented with 10 μM palmitate and 0.25 µCi/ml [U-^14^C]- or [1-^14^C]palmitic acid^52^. The ketone bodies were extracted as a non-volatile, acid-soluble product from [1-^14^C]palmitic acid incubations. Thus, after stopping the reaction with perchloric acid, 1 ml of medium was collected and transferred to a delipidated tube containing 8 volumes of chloroform/methanol (2:1, v/v) and 2 volumes of KCl (0.1 M). After shaking, it was centrifuged for 5 min at 3000 *g,* and 2 ml of the upper aqueous phase were transferred to a new tube with 4 volumes of the previous mixture, to obtain a purer sample. It was centrifuged again, and three ml aliquots of the aqueous phase were taken and transferred to vials with scintillation fluid for radioactivity determination (ketogenesis). To measure the carbon flux through the PPP pathway, cells were incubated in DMEM with 0.3 µCi/ml of [6-^14^C] or [1-^14^C]glucose^1^. Incubations were terminated after 90 min by the addition of 0.2 ml 20% perchloric acid (Merck Millipore) and, after a further 60 min, the tube containing benzethonium hydroxide (with the trapped ^14^CO_2_) was used to determine the radioactivity using a liquid scintillation analyzer. The net PPP flux was obtained by calculating the difference between the production of ^14^CO2 from glucose labeled at carbon 1 and the production of ^14^CO2 from glucose labeled at carbon 6. To measure the carbon flux from pyruvate to CO_2_ through pyruvate dehydrogenase activity, cells were incubated in DMEM with 0.2 μCi/ml [1-^14^C]pyruvate (plus 1 mM pyruvate). Incubations were terminated after 90 mins by the addition of 0.2 ml 20% perchloric acid (Merck Millipore) and, after a further 60 min, the tube containing benzethonium hydroxide (with the trapped ^14^CO_2_) was used to determine the radioactivity using a liquid scintillation analyzer. Direct pyruvate decarboxylation in the TCA cycle was measured in DMEM with 0.2 μCi/ml [2-^14^C]pyruvate (plus 1 mM pyruvate). Incubations were terminated after 90 mins by the addition of 0.2 ml 20% perchloric acid (Merck Millipore) and, after a further 60 min, the tube containing benzethonium hydroxide (with the trapped ^14^CO_2_) was used to determine the radioactivity using a liquid scintillation analyzer. In all cases, the specific radioactivity was used for the calculations. Under these experimental conditions, 70% of the produced ^14^CO_2_ and 28% of the produced ^3^H_2_O were recovered and were considered for the calculations^1^.

### Lactate determination

Lactate and glucose concentrations were measured in the culture medium spectrophotometrically^1^. For lactate concentration, a 2X reaction mixture was prepared with 0.25 M glycine, 0.5 M hydrazine, and 1 mM EDTA buffer, pH 9.5, supplemented with 1 mM NAD^+^ (N6522, Sigma) and 22.5 U lactate dehydrogenase enzyme. In a 96-well plate, 5-30 µl of medium were added per well, depending on the incubation time to allow detection of NADH, 120-145 µl of distilled water, and 150 µl of the 2X reaction mixture. Wells with medium not incubated with cells were included and used as blanks. The plate was incubated for 60-120 minutes at 37 °C, and the absorbance at 340 nm was determined. Once the absorbances of the blanks were subtracted, the lactate concentrations were calculated following the *Lambert-Beer Law*, which indicates that the absorbance is proportional to the concentration times the light path length (1 cm) and the molar extinction coefficient, the latter being 6.22 × 10^3^ M^−1^cm^−1^ for NADH and NADPH. To obtain the final concentration value, they were multiplied by the dilution factors and normalized by the mg of protein.

### Intracellular glycogen determination

Glycogen concentration was measured in primary astrocytes. First, cells were washed twice with PBS and lysed with 40 μl of 0.1 M NaOH per cm² using a scraper. The sample was then transferred to a 2 ml tube. The sample was heated at 80°C for 1 hour, cooled on ice, and then 1 ml of cold, pure ethanol was added. The sample was mixed by inversion and centrifuged at 12,000 *g* for 10 minutes. The supernatant was removed, and after allowing the pellet to dry completely, 500 µl of 50 mM sodium acetate (pH 4.8) was added. The sample was mixed vigorously, and each sample was incubated with 1 U of α-amyloglucosidase for 1 hour at 37°C. In parallel, the 2X reaction mixture described for glucose measurement was prepared, but without including the enzymes. Once the incubation was complete, the sample was mixed 1:1 with the mixture, and the initial absorbance was measured using a Uvikon XL spectrophotometer (70/99-90283 SECOMAN). The G6PDH-hexokinase enzymes were added to each cuvette and incubated for 10 minutes at RT. The absorbance measurement was repeated in the presence of the enzymes, and the values obtained in the initial measurement were subtracted from the resulting values.

### H_2_O_2_ Determination

For H_2_O_2_ assessments, AmplexRed (A12222, Thermo Fisher Scientific) was used. Cultured cells or adult brain-cell suspensions were trypsinized and incubated in KRPG buffer in the presence of 9.45 μM AmplexRed containing 0.1 U/mL horseradish peroxidase. Luminescence was recorded for 2 h at 30 min intervals using a *Varioskan Flash* (Thermo Fisher Scientific) (excitation, 571 nm; emission, 585 nm). Slopes were used for calculations of the rates of H_2_O_2_ formation.

### Mitochondrial ROS

Mitochondrial ROS were determined with the fluorescent probe MitoSox^TM^ Red (M36008, Thermo Fisher Scientific). Cultured cells were incubated with 2 μM of MitoSox for 30 min at 37 °C in a 5% CO_2_ atmosphere in Krebs-Ringer phosphate buffer (KRPG;145 mM NaCl, 4.86 mM KCl, 5.7 mM Na_2_HPO_4_, 4.09 mM NaHCO_3_, 0.54 mM CaCl_2_, 1.22 mM MgSO_4_ and 5.5 mM glucose, pH 7.35). The cells were then washed with phosphate-buffered saline (PBS; 136 mM NaCl; 2.7 mM KCl; 7.8 mM Na_2_HPO_4_·2H_2_O; 1.7 mM KH_2_PO_4_; pH 7.4) and collected by trypsinization. MitoSox fluorescence intensity was assessed by flow cytometry (FACScalibur, FL-3 channel, flow cytometer, *CellQuest^TM^* software, BD Biosciences) and expressed in arbitrary units.

### Mitochondrial membrane potential

The mitochondrial membrane potential (Δψ_m_) was assessed with MitoProbe DiIC_1_(5) (50 nM, M34151, Thermo Fisher Scientific) by flow cytometry (FACScalibur, FL-5 channel, flow cytometer, *CellQuest^TM^* software, BD Biosciences) and expressed in arbitrary units. For this purpose, cultured cells were incubated with the probe for 30 min at 37°C in KRPG buffer. Fluorescence intensities were quantified using *Paint-A-Gate™ PRO* software, and Δψ_m_ was obtained after subtraction of the potential value determined in the presence of mitochondrial uncoupler carbonyl cyanide-4-(trifluoromethoxy) phenylhydrazone (CCCP;10 µM, 15 min) for each sample, as a control of mitochondrial depolarization.

### qPCR with reverse transcription

RNA from mice hippocampus was extracted using the RNA TRI reagent according to the manufacturer’s instructions. Briefly, the brain tissue was resuspended and homogenized in 1.7 ml of TRI reagent (93289, Sigma Aldrich). Lysates were then centrifuged at 12,000 *g* for 10 minutes at 4 °C and the supernatant was collected and incubated for 15 min at RT with 170 μl of 1-bromo-3-chloropropane (B9673, Sigma Aldrich) After a 10 min 12,000 *g* centrifugation at 4 °C, the aqueous phase was collected in a new tube and 1 ml of 2-propanol (109634, Sigma Aldrich) was added. Samples were left to precipitate for 10 minutes at RT and then centrifuged at 12,000 *g* for 10 minutes at 4 °C. Supernatant was discarded, and pellets were rinsed with 1.7 ml of ethanol 75 %. Pellets were subsequently resuspended in nuclease-free H_2_O and incubated at 55 °C for 13 minutes. RNA concentration and its purity was measured using a *UV-Vis Nanodrop 2000* spectrophotometer. To evaluate the expression of the target genes, the commercial *Power SYBR® Green RNA-to-CT™ 1-Step Kit* (4389986, Applied Biosystems) was used. 100 ng of RNA were loaded in a final reaction volume mix of 20 μl, which consisted of 10 μl of *SYBR Green*, 0.16 μl of RT Enzyme, 0.6 μl of oligonucleotides 10 μM, and nuclease-free H_2_O. All reactions were performed in triplicate using the *Mastercycler ep Realplex thermocycler* (Eppendorf). The primers used were (forward and reverse, respectively) 5′-GGGAATGGTGAAGACCGTGT-3′ and 5′-CCGCATGGACTGTGGTCATGA-3′ for *cFos;* 5′-CACTCTCCCGTGAAGCCATT-3′ and 5-TCCTCCTCAGCGTCCACA-TA-3′ for *Arc,* and 5′-CAAGATCATTGCTCCTCCTG-3′ and 5′-CTGCTTGCTGATCCACATCT-3′ for *β-actin*. The relative mRNA levels were calculated using the Δ ΔΔ*C*t method, and the resulting normalized values were expressed as the fold change *versus* WT *shControl* vehicle.

### Oxygen consumption rate assessment

Oxygen consumption rates of primary astrocytes were measured in real-time in an XFe24 Extracellular Flux Analyzer (Seahorse Bioscience; *Seahorse Wave Desktop software* 2.6.1.56). This equipment measures the extracellular medium O_2_ flux changes of cells seeded in XFe24-well plates. Regular cell medium was removed and washed twice with DMEM running medium (XF assay modified supplemented with 5.5 mM glucose, 4 mM L-glutamine, 1 mM sodium pyruvate, 5 mM HEPES, pH 7.4) and incubated at 37°C without CO_2_ for 1 hour to allow cells to pre-equilibrate with the assay medium. Oligomycin, FCCP, and a mixture of rotenone and antimycin, diluted in DMEM running medium were loaded into port-A, port-B, and port-C, respectively. Final concentrations in XFe24 cell culture microplates were 2.5 μM oligomycin, 4.5 μM FCCP, 1 μM rotenone, and 2.5 μM antimycin A. The sequence of measurements was as follows. Basal level of oxygen consumption rate (OCR) was measured 3 times, and then port-A was injected and mixed for 3 min, after OCR was measured 3 times for 3 min. Same protocol with port-B and port-C. OCR was measured after each injection to determine mitochondrial or non-mitochondrial contribution to OCR. All measurements were normalized to the average three measurements of the basal (starting) level of cellular OCR of each well. Each sample was measured in 3–5 replicas. Experiments were repeated 3 times in biologically independent culture preparations. Non-mitochondrial OCR was determined by OCR after antimycin A plus rotenone injection. Maximal respiration was determined by the maximum OCR rate after FCCP injection minus non-mitochondrial OCR. ATP production was determined by the last OCR measurement before oligomycin injection minus the minimum OCR measurement after oligomycin injection. To estimate CII-sustained mitochondrial respiration were used 20 ug of mitochondria from primary astrocytes diluted in MAS buffer (70 mM sucrose, 220 mM mannitol, 5 mM KH_2_PO_4_, 5 mM MgCl_2_, 1 mM EGTA, 2 mM HEPES; pH 7.2) and from the initial respiration sustained by the addition of 5 mM succinate plus 2 μM rotenone the non-mitochondrial OCR, obtained after antimycin A injection, was subtracted.

### Cytosolic and mitochondrial ATP levels assessment

Mitochondrial (mitoATP) versus cytosolic ATP (glicoATP) production rates were determined simultaneously by *Seahorse technology* using the following injections as indicated in the figure panels at the following final concentration in the well: oligomycin (2.5 μM) was loaded into port-A and a mixture of rotenone (1µM) and antimycin A (2.5 μM) loaded into port-B. Alternatively, these ATP rates were also assessed with the single-fluorophore probe iATP-GFP^53^. This probe consists of the GFP protein fused to the F_0_F_1_-ATPase from Bacillus PS3. iATP-GFP responds to ATP concentrations in the range of 30 μM to 3 mM, with fast changes in fluorescence emissions. iATP-GFP probe expression plasmid was acquired from *Addgene* (*Plasmid #102553*). Since the expression of the iATP-GFP probe was governed by a GFAP promoter, its sequence was subcloned into a pcDNA3.1(+) plasmid, which possesses the strong and ubiquitous CMV promoter. Astrocytes were seeded on 8-well IBIDI plates. The day before the assay, cells were transfected with the plasmid containing the iATP-GFP probe using the *Lipofectamine™ LTX with Plus™ Reagent* following the manufactureŕs instructions. The day of the assay, media was replaced by 200 μl of KRPG solution. Cells were observed in the *Andor Dragonfly spinning disk* confocal microscopy (AndorTechnology, Oxord Instruments). Images were obtained with the Cy2 channel (λEx = 488 nm/λEm = 521 nm), with a laser potency of 10 % and an exposure time of 700 ms. Each image was obtained covering the entire cell thickness with 1 μm steps. One image of the initial conditions was captured. Afterwards, KRPG was replaced by KRP, which presents the same composition but lacks glucose. KRP was also supplemented with 2-deoxyglucose 10 mM (D6134, Sigma Aldrich) or oligomycin 2.5 μM to determine cito or mitoATP, respectively. Micrographs were obtained for 10 minutes every 30 seconds, and image quantification was performed with the *FIJI software*. To assess fluorescence loss over time, intensity values of each time point were plotted, and the slope of the linear regions (R2 > 0.9) was analyzed.

### ATP hydrolysis in previously frozen samples (HyFS)

ATP hydrolysis capacity or State 4 acidification rates were measured using Agilent Seahorse XFe24 as described^54^. Briefly, plates were loaded with 40 μg of cell lysate or 20 μg of mitochondria in MAS buffer. Cell lysates were prepared by subjecting the samples to 4 cycles of freeze-thaw (liquid nitrogen-37°C water bath) before measuring protein concentration. Initial respiration of the samples was sustained by the addition of 5 mM succinate plus 2 μM rotenone in MAS after centrifugation. Injections were performed as indicated in the figure panels at the following final concentration in the well: Antimycin A (AA) (2 μM), oligomycin (5 μM), FCCP (1 μM), ATP (20 mM). To assess maximal ATP concentration, ATP was injected consecutively. ATP hydrolysis was determined by the maximum acidification rate (ECAR) after ATP/FCCP injection minus the minimum ECAR measurement after oligomycin injection.

### Total ATP levels determination

To determine total ATP levels in primary cultures of astrocytes, a commercial bioluminescence kit (A22066, Thermo Fisher) following the manufactureŕs instructions was used. Opaque 96-well plates were used, and luminescence was evaluated in a *Variskan Flash luminometer* (Thermo Fisher) at λ=560 nm every 5 minutes for 20 minutes at 28 °C. In parallel, a standard curve was run with ATP standards (from 0 to 7.5 μM), from which the concentration of the samples was extrapolated.

### NAD^+^ and NADH(H^+^) determinations

To determine NAD^+^/NADH(H^+^) levels, a commercial kit (MAK460, Sigma) was used following the manufacturer’s instructions. Excitation intensities of 530 nm and emission at 585 nm were determined. The 96-well plate was incubated for 10 minutes in the dark at RT, and the measurement was carried out at time 0 and after 10 minutes. To calculate the NAD^+^ or NADH(H^+^) concentration, a standard line was run in parallel from 0 (blank) to 1 μM NAD^+^ or NADH(H^+^) from which the sample concentrations were extrapolated.

### Protein determinations

Protein samples were quantified by the BCA protein assay kit (Thermo Fisher) using BSA as a standard.

### Western Blotting

Cells were lysed in RIPA buffer (1% sodium dodecylsulfate, 10 mM ethylenediaminetetraacetic acid (EDTA), 1 % (vol/vol) Triton X-100, 150 mM NaCl and 10 mM Na_2_HPO_4_, pH 7.0), supplemented with protease inhibitor mixture (Sigma), 100 μM phenylmethylsulfonyl fluoride, and phosphatase inhibitors (1 mM o-vanadate). Samples were boiled for 5 min. Aliquots of cell lysates (40-60 μg of protein) were subjected to SDS/PAGE on an 8 to 15% (vol/vol) acrylamide gel (MiniProtean; Bio-Rad), including *PageRuler Prestained Protein Ladder* (Thermo). The resolved proteins were transferred electrophoretically to nitrocellulose membranes (0.2 µm, BioRad) at 60 V for 90 minutes. Membranes were blocked with 5% (wt/vol) low-fat milk (Sveltesse, Nestle) in TTBS (20 mM Tris, 150 mM NaCl, and 0.1% (vol/vol) Tween 20, pH 7.5) for 1 h. After blocking, membranes were immunoblotted with primary antibodies overnight at 4 °C. After incubation with horseradish peroxidase-conjugated goat anti-rabbit IgG (1/20,000, sc-2020, Santa Cruz Biotechnologies), goat anti-mouse IgG (1/30,000, 1858413, Pierce), and rabbit anti-goat IgG (1/20,000, sc-2701, Santa Cruz Biotechnologies), membranes were immediately incubated with the enhanced chemiluminescence kit *WesternBright ECL* (Advansta), or *SuperSignal West Femto* (Thermo) before exposure to Fuji Medical X-Ray film (Fujifilm) or *Fusion FX Vilber transilluminator*, and the autoradiograms were scanned. At least three biologically independent replicates were always performed, although only one representative Western blot is shown in the main figures. The protein abundances of all Western blots per condition were measured by densitometry of the bands on the films using *ImageJ 1.48u4 software* (National Institutes of Health). They were normalized by loading control protein. The resulting values were used for statistical analysis. Uncropped scans of Western blots replicas are shown in the *Source Data file*.

### Primary antibodies for Western blotting

Immunoblotting was performed with anti-PFKFB1,4 (1/500) (sc-10096; Santa Cruz Biotechnologies), anti-PFKFB2 (1/500) (ab234865; Abcam), anti-Pfkfb3 (1/1,000) (H00005209-M08; Novus Biologicals), anti-Pfkfb3 (1/1,000) (ab181861; Abcam), anti-PDHA1 (1/1,000) (#3205; Cell Signaling), anti-phospho-Ser293-PDHA1 (1/1,000) (#31866; Cell Signaling), anti-MCT2 (1/500) (sc-166925; Santa Cruz Biotechnologies), anti-NDUFA9 (1/1,000) (ab14713; Abcam), anti-UQCRC2 (1/1,000) (ab14745; Abcam), anti-GFAP (1/500) (G6171; Sigma), anti-MAP2 (1/500) (ab11268; Abcam), anti-IBA1 (1/500) (019-19741; Wako) and anti-β-actin (1/30,000) (A5441; Sigma).

### Mitochondrial isolation

To obtain the mitochondrial fraction, cell pellets were frozen at −80 °C and homogenized (ten-twelve strokes) in a glass-Teflon Potter–Elvehjem homogenizer in buffer A (83 mM sucrose and 10 mM MOPS; pH 7.2). The same volume of buffer B (250 mM sucrose and 30 mM MOPS) was added to the sample, and the homogenate was centrifuged (1,000 *g*, 5 min) to remove unbroken cells and nuclei. Centrifugation of the supernatant was then performed (12,000 *g*, 3 min) to obtain the mitochondrial fraction, which was washed in buffer C (320 mM sucrose; 1 mM EDTA and 10 mM Tris-HCl; pH 7.4)^29^. Mitochondria were suspended in buffer D (1 M 6-aminohexanoic acid and 50 mM Bis-Tris-HCl, pH 7.0) for blue native gel electrophoresis.

### Blue native gel electrophoresis

For the assessment of complex I organization, digitonin-solubilized (4 g/g) mitochondria (10–20 μg) were loaded in NativePAGE Novex 3–12% (vol/vol) gels (Life Technologies). The electrophoresis was carried out at 40 V overnight. Next, a direct electrotransfer was performed, followed by immunoblotting against mitochondrial complex I antibodies NDUFA9 (1/1,000) (ab14713; Abcam), complex III antibody UQCRC2 (1/1,000) (ab14745; Abcam). Direct transfer of BNGE was performed after soaking the gels for 20 min (4 °C) in carbonate buffer (10 mM NaHCO_3_; 3 mM Na_2_CO_3_·10H_2_O; pH 9.5–10). Proteins were transferred to polyvinylidene fluoride (PVDF) or 0.2 μm nitrocellulose membranes (Hybond^®^, Amersham Biosciences) and were carried out at 300 mA, 60 V, 1.5 h at 4 °C in carbonate buffer.

### Mouse perfusion, immunohistochemistry, and image analysis

Animals were deeply anesthetized by intraperitoneal injection of a mixture (1:3) of xylazine hydrochloride (Rompum, Bayer) and ketamine hydrochloride/chlorbutol (Imalgene; Merial) using 1 ml of the mixture per kg of body weight, and then perfused intraaortically with 0.9 % NaCl followed by 5 mL/g body weight of fixative solution [4 % (wt/vol) paraformaldehyde, depolymerized with NaOH and stabilized with 10 % (v/v) methanol, in 0.1 M phosphate buffer, pH 7.4]. After perfusion, brains were dissected out sagittally into two parts and postfixed, using fixative solution, for 24 h at 4 °C. Afterwards, brains were cryoprotected in 30 % (wt/vol) sucrose in PBS. After cryoprotection, 10, 20, and 40-μm-thick sagittal sections were obtained with a cryostat (Leica; CM1950 AgProtect). Sections were then incubated with primary antibodies: i)1/500 anti-MAP2 (AP-20; abcam 11268), 1/500 anti-GFAP (G6171; Sigma), 1/500 anti-Iba1 (Wako, 019-19741), in in 0.2 % Triton X-100 (Sigma-Aldrich) and 10 % goat serum (Jackson ImmunoReseach) in 0.1 M PBS for 72 h at 4°C. Later, fluorophore-conjugated secondary antibodies ii) mouse Cy^TM^2, 115-225-003; rabbit Cy^TM^3, 111-165-003, 1/1000 (Jackson ImmunoResearch) in 0.05 % Triton X-100 and 2 % goat serum in 0.1 M PB for 2 h at room temperature^55^. For nuclear staining, sections were incubated with DAPI 0.1 μg/ml in PB for 10 min to stain nuclei. Finally, sections were mounted with Fluoromont (Sigma) aqueous mounting medium. Confocal images were taken with a scanning laser confocal microscope (*Spinning Disk ANDOR DragonFly Nikon Ti2-E*) with three lasers, 405, 488, and 561 nm, and equipped with *Fusion 2.2* acquisition software. All images acquired by confocal microscopy were processed and, after deconvolution, analyzed using the free software *FIJI* (ImageJ64). The quantification of the different parameters analyzed was performed on *Z-Project* micrographs of three different regions of the CA1 from each hippocampus, taken from three serial sagittal sections per mouse. Both the percentage of area occupied by MAP2, GFAP, and IBA1, as well as the mean fluorescence intensity, were measured after converting the images to 8-bit and performing a *Huang threshold*. To characterize cell roundness in astrocytes and microglia, a particle analysis plugin was applied after the *threshold*.

### Metabolomics analysis

The most medial portion of the right hemisphere (approximately 70 mg) of 1-year-old WT or astrocyte-specific *Pfkfb3*-KO mice (n=6 for each condition) was snap frozen in liquid nitrogen and used for untargeted metabolomics analysis (Metabolon Incorporated^®^). Raw data were extracted, peak-identified, quality-control processed, curated, normalized, and log2 transformed by the Metabolon service. Peaks were quantified with the area under the curve. For studies spanning multiple days, a data normalization step was performed to correct variation resulting from instrument inter-day tuning differences. Essentially, each compound was corrected in run-day blocks by registering the medians to equal one (1.00) and normalizing each data point proportionately. Following imputation of missing values with the minimum observed value for each compound, Welch’s two-sample *t* tests were used to identify compounds significantly different between experimental groups. Using these filters, we accepted a high estimate of the false discovery rate (q value ≤ 0.05), although other considerations were implemented to determine whether a result merited further scrutiny. Such evidence included (i) biological relevance given the genetic background context; (ii) inclusion in a common pathway with a highly significant compound; (iii) residing in a similar functional biochemical family with other significant compounds; or (iv) correlation with other experimental approaches. Graphs corresponding to statistical analysis were carried out with *GraphPad 8.0* and the online tool *MetaboAnalyst 5.0*.

### Behavioral tests

Male and female mice (6 weeks post AAV injections, i.e., 4-5 months old) were left to acclimatize in the room for not less than 1 hour at the same time slot of the day (9 am-14 pm). Both sexes were evaluated separately. Tracking was carried out once at a time and carefully cleaning the apparatus with 70% ethanol between trials to remove any odor cues. An ANY-box^®^ core (ANY-Maze, Stoelting Europe) was used, which contained a light grey base and an adjustable perpendicular stick holding a camera and an infrared photo-beam array connected to an AMi-maze^®^ interface to track the animal movement and to detect rearing behavior, respectively. For the *Openfield* test, a 40 cm x 40 cm x 35 cm (w, d, h) black infrared transparent Perspex insert was used, and the arena was divided into three zones, namely border (8 cm wide), center (16 % of total arena), and intermediate (the remaining area). The test lasted for 10 min, and the distance travelled, the number and duration of rearings, the immobile time, and the time spent in each zone were measured. As a variant of the *Openfield* test, the *Hole Board test* was used, adding a metal structure with 16 holes evenly distributed across the surface. The distance of the infrared sensors was adjusted to detect the mousés explorations, which consisted of inserting its head into one of the holes. The animal was placed in the box and allowed to explore for 5 minutes. The number of explorations was subsequently analyzed. To evaluate fear and anxiety levels, we used *Black & White Box.* This test consisted of placing the animal in two rooms: one bright with transparent walls, where it feels more exposed and stressed, and a darker one with opaque walls, where it feels more protected. To perform this test, a Plexiglas cubicle was placed with an internal separation between the two compartments, one opaque and the other transparent, with a passageway to allow the animal to change rooms. The animal was placed in the transparent part and allowed to explore for 10 minutes. The time and distance traveled in both rooms were evaluated. The *Rotarod test* (Rotarod apparatus, Model 47600, Ugo Basile) was used to analyze motor balance and coordination. Mice were previously trained for three consecutive days, two days before the test. The rotarod conditions were a gradual acceleration from 4 to 40 r.p.m., reaching the final speed at 270 sec. The day of the test, mice latency to fall from the rotarod was recorded. To analyze the short-term memory, we used the *Novel Object Recognition test* (Stoelting). Mice were accustomed to the ANY-box^®^ core for 10 min, 24 hours before the test. Mice were left to explore two identical equidistant objects (familiar object) for 5 min (the familiarization phase) and returned for 30 min into its cage. One object was replaced by another of similar size (novel object), and mice were returned to the arena to explore the objects for another 5 min (the test phase). The software scored the investigation of the objects when the head of the animal was looking at the object within a distance of 20 mm. The ability to recognize the novel object was determined as discrimination index (DI) calculated as [DI = (T_N_ – T_F_) / (T_N_ + T_F_)], where T_N_ is the time spent exploring the new object and T_F_ is the time spent exploring the familiar object. For the *Barnes maze test*, we used equipment consisting of a grey circular platform (90 cm in diameter, elevated 90 cm above the floor). Along its perimeter, there were 20 evenly spaced holes. The maze has one removable escape box that could be fitted under any of the holes and was filled with the animal bedding before each experiment. Black and white patterned pictures were used as spatial visual cues. All sessions were performed under a room lighting of 400 lux to increase the mouse aversion for the platform. The test consisted of three phases. First, the habituation phase, where the animals were left to explore the platform freely for 5 minutes, 1-day before the training sessions. Afterwards, the animals underwent the training phase where they were allowed to locate the escape hole for a maximum of 5 minutes, for 4 days with 3 sessions per day. Finally, for the probe phase, mice were tested for spatial memory. In this session, the escape box was removed, and the platform was virtually divided into four quadrants, each containing five holes. Mice were allowed to explore the maze for 5 minutes, and the time spent in the quadrant that previously contained the escape box was quantified to evaluate spatial memory and learning^56^. For the *Y-maze test* (working memory test), we used a maze composed of three arms in the form of a Y shape labeled A, B, and C, respectively. Each arm is 30 cm long, 5 cm wide, and 12 cm high. Animals were placed at the end of arm A and allowed to move freely through the maze for 5 minutes^10^. The ANY-maze software considered an arm entry was counted when 90% of the mousés body was inside the arm. Correct spontaneous alternation was defined as the number of sequential entry triads in three different arms (ABC, ACB, BAC, BCA, CAB, CBA). The percentage of correct spontaneous alternations was calculated as follows: [SA = (T_S_/(T_AE_-2)*100], where T_S_ is the total number of correct spontaneous alternations and T_AE_ is the total number of arm entries. The number of evaluated animals per test is specified in the figure legends.

### *cFos* and *Arc* determination

Animals were introduced in an ANY-box^®^ core with red and blue patterns in the walls to stimulate them and were allowed to freely explore the box for 15 min. Then mice were returned to their cages for 90 minutes, a time at which *cFos* and *Arc* expression reach their peak after stimulation^57,58^. Afterwards, the animals were sacrificed by cervical dislocation, their brains were removed, and the prefrontal cortex and hippocampus were flash frozen in liquid nitrogen and stored at -80°C before RNA extraction.

### Statistical Analysis

For simple comparisons, we used unpaired two-tailed Student’s *t-test* or unpaired Mann-Whitney test when the data do not follow a normal distribution or the variances between populations are different. For other multiple-values comparisons, we used one-way or two-way ANOVA followed by Tukey or Dunnett post-hoc tests, depending on whether we want to compare all the conditions with each other or with a specific control, to minimize familywise error rate, which is the probability of making at least one type I error (false positive) when conducting multiple comparison. In the metabolomics study, the Welch *t-test* was chosen, following the recommendation of Metabolon. This test is parametric, but, unlike the Student’s *t-test*, it does not assume homogeneity of variances. This choice is based on the company’s day-to-day experience in metabolomics data processing, providing a more robust statistical method. In all cases, a *p-value* ≤ 0.05 was considered statistically significant, although the *p-values* are indicated numerically in the graphs. All tests used are indicated in each figure legend. The statistical analysis was performed using the *GraphPad Prism v8 software*. The number of biologically independent culture preparations or animals used per experiment is indicated in the figure legends.

## Reporting summary

Further information on research design is available in the Nature Portfolio Reporting Summary linked to this article.

## Data availability

Source data are provided with this paper.

---

**Extended Data Fig. 1.**
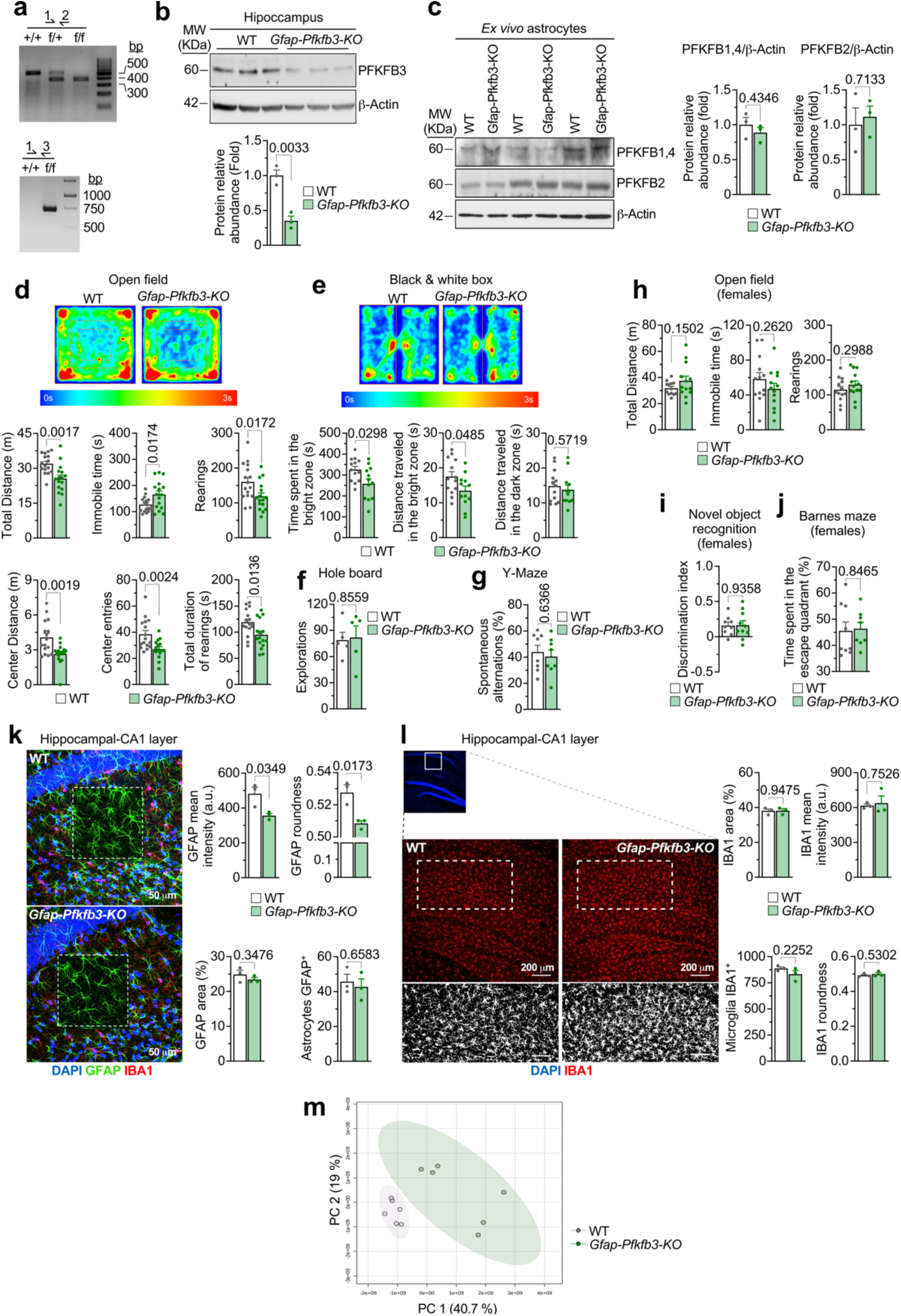
Confirmation of astrocyte-specific Pfkfb3 knockout and assessment of general behavior and glial morphology. **(a)** Genotyping analysis confirming Pfkfb3 alleles. *Upper gel*: polymerase-chain reaction (PCR) amplification products for wild-type (+/+), heterozygous (f/+), and homozygous (f/f) genotypes using primer pair 1 and 2 (expected band sizes indicated). *Lower gel*: PCR with primer pair 1 and 3 differentiating wild-type (+/+) and knockout (f/f) alleles. Related to Fig. 1a. **(b)** Validation of Pfkfb3 protein reduction in hippocampus: *Top*: Western blot for PFKFB3 and β-Actin in WT and *Gfap-Pfkfb3-KO* hippocampal samples. *Bottom*: Quantification of relative PFKFB3 abundance (fold change normalized to WT). Data are mean ± S.E.M. *P* value is indicated; n=3 mice per genotype; Unpaired Student’s t-test, two-tailed. Uncropped western blot replicas are shown as Source Data file. Related to Fig. 1b. **(c)** PFKFB1,4 and PFKFB2 isoforms protein abundances in astrocytes immunomagnetically isolated from mouse brain. *Left*: PFKFB1,4, PFKFB2, and β-Actin in WT and *Gfap-Pfkfb3-KO* astrocytes. *Right*: Quantification of the bands. Data are mean ± S.E.M. *P* values are indicated; n=3 mice per genotype; Unpaired Student’s t-test, two-tailed. Uncropped western blot replicas are shown as Source Data file. Related to Fig. 1b. **(d)** Open field behavioral test. *Top*: representative heat maps of locomotor activity in WT and *Gfap-Pfkfb3-KO* male mice. *Bottom*: quantifications of the indicated parameters. Data are mean ± S.E.M. *P* values are indicated; n=16 mice per genotype; Unpaired Student’s t-test, two-tailed. Related to Fig. 1d-f. **(e)** Anxiety-like behavior assessed by black-and-white box. *Top*: heatmaps for WT and *Gfap-Pfkfb3-KO* male mice. *Bottom*: quantifications of the indicated parameters. Data are mean ± S.E.M. *P* values are indicated; n=12 (time spent in the bright zone WT, distance travelled in the bright or dark zones *Gfap-Pfkfb3-KO*) or 13 (time spent in the bright zone *Gfap-Pfkfb3-KO*, distance travelled in the bright or dark zones WT) mice; Unpaired Student’s t-test, two-tailed. Related to Fig. 1d-f. **(f)** Exploratory behavior in hole board test in WT and *Gfap-Pfkfb3-KO* male mice. Data are mean ± S.E.M. *P* value is indicated; n=5 mice per genotype; Unpaired Student’s t-test, two-tailed. Related to Fig. 1d-f. **(g)** Y-maze spontaneous alternation as a measure of working memory in WT and *Gfap-Pfkfb3-KO* male mice. Data are mean ± S.E.M. *P* value is indicated; n=8 mice per genotype; Unpaired Student’s t-test, two-tailed. Related to Fig. 1d-f. **(h)** Open field behavioral parameters in WT and *Gfap-Pfkfb3-KO* female mice. Data are mean ± S.E.M. *P* values are indicated; n=13 mice per genotype; Unpaired Student’s t-test, two-tailed. Related to Fig. 1d-f. **(i)** Novel Object Recognition analysis in WT and *Gfap-Pfkfb3-KO* female mice. Data are mean ± S.E.M. *P* value is indicated; n=9 (WT) or 10 (*Gfap-Pfkfb3-KO*) mice; Unpaired Student’s t-test, two-tailed. Related to Fig. 1d-f. **(j)** Barnes maze spatial memory analysis in WT and *Gfap-Pfkfb3-KO* female mice. Data are mean ± S.E.M. *P* value is indicated; n=8 (*Gfap-Pfkfb3-KO*) or 9 (WT) mice; Unpaired Student’s t-test, two-tailed. Related to Fig. 1d-f. **(k)** Analysis of the astrocyte morphology in the hippocampal CA1 layer in WT and *Gfap-Pfkfb3-KO* mice. *Left*: immunofluorescence images stained for DAPI (blue), GFAP (green), and IBA1 (red), with dashed boxes marking analyzed regions. Scale bar: 50 µm. *Right panels*: quantification of the indicated parameters. Data are mean ± S.E.M. *P* values are indicated; n=3 mice per genotype; Unpaired Student’s t-test, two-tailed. Related to Fig. 1g. **(l)** Microglial analysis in the hippocampal CA1 layer of WT and *Gfap-Pfkfb3-KO* mice. *Left*: DAPI and IBA1 images (scale bar 200 µm), with magnified view below. *Right panels*: quantification of the indicated parameters. Data are mean ± S.E.M. *P* values are indicated; n=3 mice per genotype; Unpaired Student’s t-test, two-tailed. Related to Fig. 1g. **(m)** Principal component analysis (PCA) plot showing clustering of WT (gray) and *Gfap-Pfkfb3-KO* (green) samples according to principal components PC1 (40.7%) and PC2 (19%). N=6 mice per genotype. Related to Fig. 1h.

**Extended Data Fig. 2.**
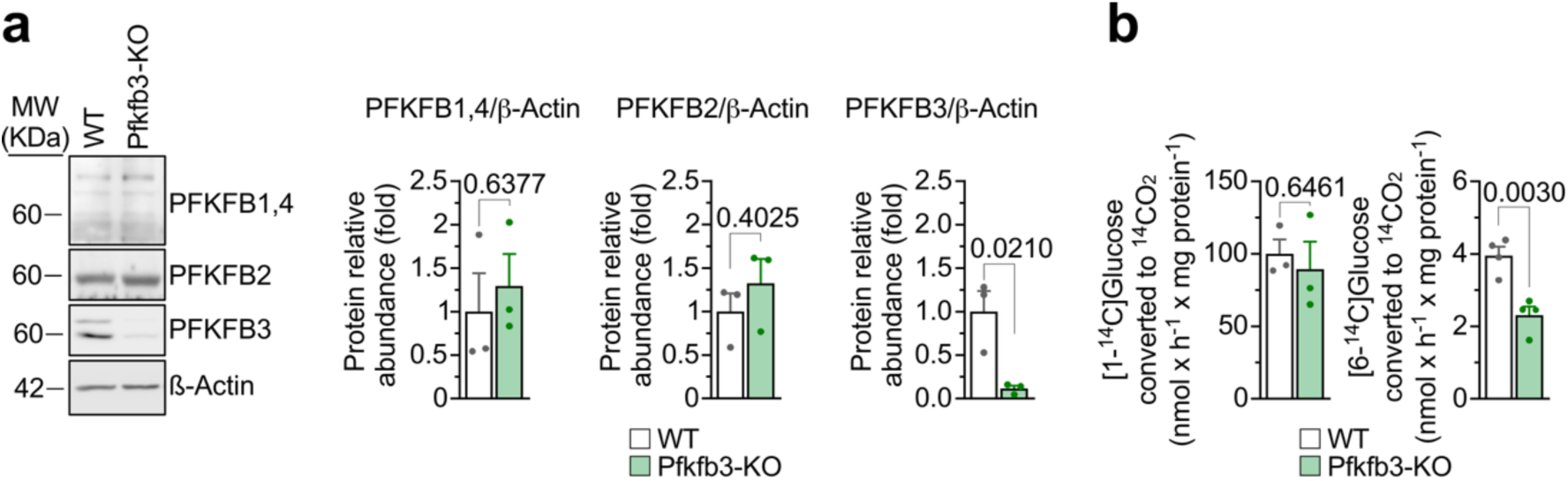
PFKFB isoforms and alternate glucose metabolism in Pfkfb3-KO astrocytes. **(a)** Expression of PFKFB isoforms in WT and Pfkfb3-KO primary astrocytes. Western blots (left) and their quantification (*right*) are shown. Data are mean ± S.E.M. *P* values are indicated; n=3 biologically independent samples per genotype; Unpaired Student’s t-test, two-tailed. Related to Fig. 2b. **(b)** Glucose metabolic fluxes of [1-^14^C]glucose (via pentose-phosphate pathway) and [6-^14^C]glucose (via glycolysis/TCA) to ^14^CO_2_, in WT and Pfkfb3-KO primary astrocytes. Data are mean ± S.E.M. *P* values are indicated; n=3 (1-^14^C) or 4 (6-^14^C) biologically independent samples per genotype; Unpaired Student’s t-test, two-tailed. Related to Fig. 2g.

**Extended Data Fig. 3.**
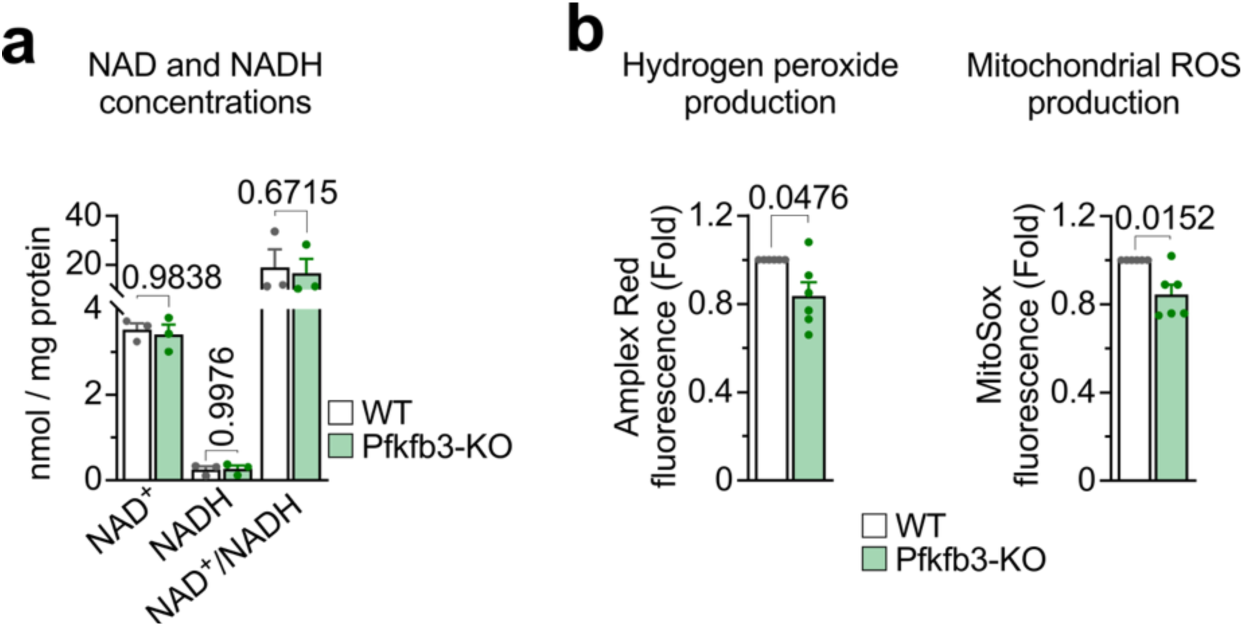
Redox parameters in Pfkfb3-KO astrocytes. **(a)** NAD⁺, NADH, and NAD⁺/NADH ratio in WT and Pfkfb3-KO primary astrocytes. Data are mean ± S.E.M. *P* value is indicated; n=3 biologically independent samples per genotype; Unpaired t test, two-tailed. Related to Fig. 3. **(b)** Hydrogen peroxide (H_2_O_2_) and mitochondrial reactive oxygen species (ROS) production in WT and Pfkfb3-KO primary astrocytes. Data are mean ± S.E.M. *P* value is indicated; n=6 biologically independent samples per genotype; Unpaired Mann-Whitney test, two-tailed. Related to Fig. 3.

**Extended Data Fig. 4.**
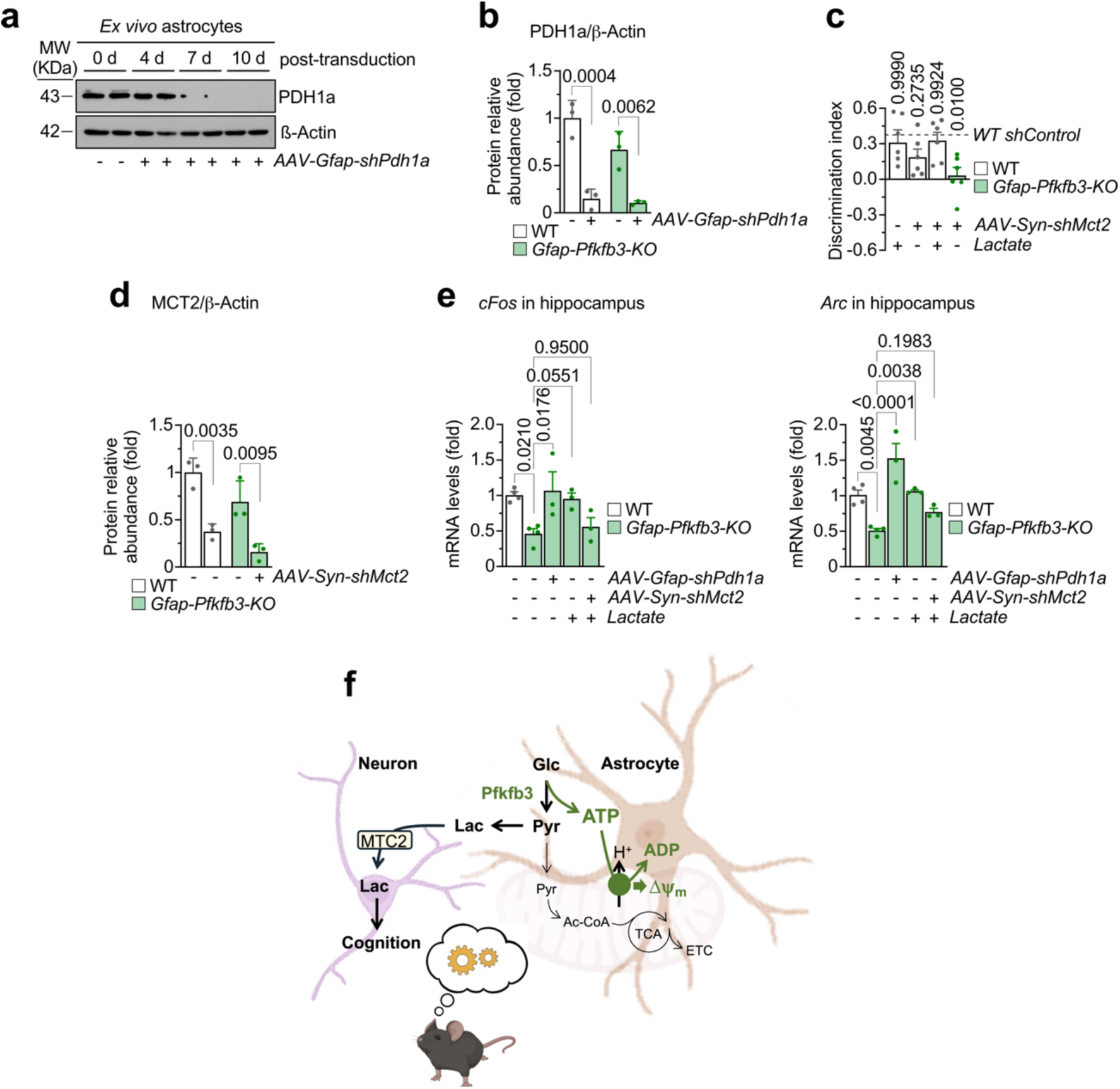
Control experiments in astrocyte-specific *Pdh1a* and *Mct2 in vivo* silencing and lactate administration and modulation of neuronal activity markers. **(a)** Time course of PDH1a protein knockdown in immunomagnetically isolated astrocytes following *in vivo AAV-Gfap-shPdh1a* transduction in WT mice. For the rest of the experiments, 10 days post-transduction was used, when PDH1a protein was absent according to this representative Western blot. A biologically independent replica was performed. ß-Actin was used as loading control. Related to Fig. 4d. **(b)** Quantification of the Western blots against PDH1a in immunomagnetically isolated astrocytes from WT and *Gfap-Pfkfb3-KO* mice 10 days after transduction with *AAV-Gfap-shPdh1a.* Data are mean ± S.E.M. *P* values are indicated; n=3 mice per genotype and treatment; two-way ANOVA followed by Tukey. Related to Fig. 4d. **(c)** Control experiments for the discrimination index quantification in the Novel object recognition test in WT and *Gfap-Pfkfb3-KO* male mice treated or not with lactate. The dashed horizontal line shows the discrimination index value in the WT *shControl* condition, that is the value shown in Fig. 4e, *right panel*, condition “WT – *AAV-Syn-shMct2* – lactate”. Data are mean ± S.E.M. *P* values are indicated; n=6 mice per genotype and treatment; two-way ANOVA followed by Dunnet. Related to Fig. 4e. **(d)** Quantification of the Western blots against MCT2 in immunomagnetically isolated neurons from WT and *Gfap-Pfkfb3-KO* mice 10 days after transduction with *AAV-Syn-shMct2.* Data are mean ± S.E.M. *P* values are indicated; n=3 mice per genotype and treatment; two-way ANOVA followed by Tukey. Related to Fig. 4e. **(e)** Activity-dependent neuronal gene expression in hippocampus, as judged by *cFos* (*left*) and *Arc* (*right*) mRNA abundances in WT and *Gfap-Pfkfb3-KO* male mice treated or not with *AAV-Gfap-shPdh1a*, *AAV-Syn-shMct2* or lactate. Data are mean ± S.E.M. *P* values are indicated; n=3 (Gfap-Pfkfb3-KO except no treatments in *cFos*) or 4 (WT and *Gfap-Pfkfb3-KO* in no additions in *cFos*) mice per genotype and treatment; one-way ANOVA followed by Dunnet. Related to Fig. 4d,e. **(f)** Graphical summary describing the main message of this work. We show that in astrocytes, Pfkfb3 is instrumental in sustaining glycolytically generated ATP, which is hydrolyzed in mitochondria by the ATP synthase in its reverse activity mode to maintain the inner mitochondrial membrane potential (Δψ_m_). In this way, glycolytically derived pyruvate is not necessary to provide acetyl-CoA to sustain TCA cycle and ETC activities, and therefore it is converted into lactate, which is taken up by neurons to sustain function and cognition.

## Acknowledgements

We acknowledge the technical assistance of M. Resch, L. Martin and E. Prieto-Garcia. E. Prieto-García was a recipient of a position by the Fondo Social Europeo, Youth Employment Initiative, Junta de Castilla y León. JPB is funded by MICIU/AEI (PID2022-138813OB-I00 /10.13039/501100011033 and FEDER, UE), la Caixa Foundation (grant agreement LCF/PR/HR23/52430016), the European Union’s Horizon Europe research and innovation program under the MSCA Doctoral Networks 2021 (101072759; FuEl ThEbRaiN In healtThY aging and age-related diseases, ETERNITY, and the European Research Council (ERC) Advanced Grant NeuroSTARS (ref. 101199747). AAP is funded by the Instituto de Salud Carlos III (PI24/00810, PMP22/00084 and RD24/0009/0005 co-funded by the European Union) FEDER; Junta de Castilla y León (CSI011P23).

## Author contributions

Conceived the idea and designed research: J.P.B.

Performed research: P.A.-B., D.J.-B., J.A., R.L., M.A.-D., S.Y.-S., B.M.-F., D.G.-R., R.A.-P. V.B.J., L.S.-O., R.M.-F., S.G., S.G.-G.

Analysed data: J.P.B., A.A., P.A.-B., D.J.-B., E.F., J.A.E.

Contributed materials: P.C., G.B. Wrote the manuscript: J.P.B.

Edited and approved the manuscript: all co-authors.

